# Host microbiome determines host specificity of the human whipworm, *Trichuris trichiura*

**DOI:** 10.1101/2025.05.09.653168

**Authors:** Klara Aliz Stark, Tapoka T. Mkandawire, Charlotte Tolley, Seona Thompson, Rachael Fortune-Grant, Cordelia Brandt, Simon Clare, Trevor D. Lawley, Cinzia Cantacessi, Alexandre Almeida, Richard K. Grencis, Peter Nejsum, Matthew Berriman, María A. Duque-Correa

## Abstract

Long-term whipworm-host co-evolution has resulted in tropism for the caecum of specific hosts, an organ with the densest microbial population in the body. Here, we demonstrate that the host specificity of human whipworms *(Trichuris trichiura)* is host microbiome-driven. We successfully establish a *T. trichiura* infection in a non-primate host using a humanised-microbiota mouse model. We further show, *in vitro*, that hatching of *T. trichiura* was triggered by mucosal scrapings of the caecum of human microbiota-associated mice, but not from wild-type mice, which only induced *T. muris* hatching. Comparative metagenomic analysis of the murine versus humanised microbiomes directly implicated specific bacterial species in *T. trichiura* egg hatching. Additionally, we demonstrate that host tissue does not directly determine host specificity, as *T. trichiura* readily infected mouse caecaloids. Our findings indicate that host-microbiome-whipworm co-evolution has resulted in exquisite bacterial-whipworm egg interactions critical for hatching and development of these parasites in their definitive hosts.

## Introduction

Human-infective whipworms (*Trichuris trichiura*) are soil-transmitted helminths that have been associated with humans for millennia, currently infecting ∼500 million people worldwide ^1–4^. Infections can cause Trichuriasis, a major neglected tropical disease leading to high chronic morbidities with dire socio-economic consequences in affected countries ^5,6^. No vaccines are available, and control relies on administration of benzimidazole anthelmintics that fail to clear infections effectively ^7,8^. Novel, effective and sustainable tools to tackle whipworm infections are therefore urgently needed.

Together with the human whipworm, at least another 70 *Trichuris* species infect wild and domestic mammals ^6,9^. Infections begin by ingesting whipworm eggs present in contaminated food, water and soil, which hatch upon arrival in the large intestine. All *Trichuris* species show the same tissue tropism, inhabiting the epithelium of the caecum and proximal colon of their definitive hosts, a niche within the gastrointestinal tract (GIT) with the densest microbial population ^1,10^. Whipworms have evolved closely with their host and its complex intestinal microbiota for millions of years to successfully exist as long-lived, chronic infections ^10,11^. Long-term co-adaptive processes may have led to the development of the high degree of host specificity of *Trichuris* species. Indeed, *T. trichiura,* an obligate parasite of human and other primates ^2,12,13^, is unable to establish a patent infection in non-primate experimental animals such as mice or pigs, which are naturally infected by *T. muris* and *T. suis*, respectively, and serve as models for human whipworm infection biology and pathogenesis ^6,9,14^.

*Trichuris* host specificity could be determined by: (1) unique, yet to be identified, host intestinal epithelia cues, which potentially include components of the mucus layer or cell membrane receptors, and mediate parasite tissue recognition, invasion and colonisation, or (2) interactions with specific members of the host microbiota. Studies on the mouse and pig whipworms indicate that microbial communities from the GIT of their host are critical for whipworm hatching, establishment and persistence as chronic infections ^10^. Specifically, *T. muris* is unable to hatch and infect mice in the absence of microbiota (germ-free mice) ^11^, and infectivity is restored upon colonisation with a faecal transplant from uninfected mice or monocolonisation with *Bacteroides thetaiotaomicron* ^11^, *Escherichia coli* ^15^, *Staphylococcus aureus* ^16^ or *Paraclostridium sordellii* ^17^. Consistently, *T. muris* eggs hatch *in vitro* in response to mouse caecum explants ^18^ and murine luminal caecal and colon contents ^11,19^, and monocultures of specific Gram-negative and Gram-positive bacteria ^15–18,20–23^. In contrast, these bacteria do not trigger *T. suis* hatching *in vitro* that can only be induced by exposure to mucosal scrapings of the pig ileum, caecum and colon ^24,25^. Intriguingly, while bacterial isolates from the faeces of *T. trichiura*-infected individuals, specifically the obligate anaerobe Clostridiales species *Paraclostridium sordellii* and *Romboutsia hominis,* induce hatching of *T. muris,* only *P. sordellii* trigger low levels of *T. trichiura* hatching after six days of co-culture ^17^, a time that surpasses the normal gut transit time (≈ 28 h) ^26^. Thus, the members of the human intestinal microbiota and the conditions required to induce hatching of *T. trichiura* are currently poorly understood. We hypothesise that host specificity in *Trichuris* is determined by the host gut microbiome, with each whipworm species having evolved to hatch, develop and persist in response to specific interactions with bacteria associated with the mucosa of the caecum and proximal colon of their respective hosts.

In this study, we investigated the role of human intestinal microbiota on the hatching and host specificity of *T. trichiura* using human microbiota-associated (HMA) mice ^27–29^. HMA mice successfully sustained infections with *T. trichiura*, but not with *T. muris*, and mucosal scrapings, but not luminal contents, from the caecum of HMA mice induced *in vitro* hatching of *T. trichiura*. Conversely, wild-type (WT) mice harbouring murine intestinal microbiota supported infections with *T. muris,* and their caecal mucosal scrapings and luminal contents triggered the hatching of *T. muris*, but not of *T. trichiura*. Hatching of both *Trichuris* species was observed under both aerobic and anaerobic conditions and was shown to be mediated by proteases. Comparative metagenomic analyses of the caecal microbiome of WT and HMA mice led to the identification of bacterial species implicated in *T. trichiura* egg hatching, which was triggered *in vitro* by several anaerobes of the human gut microbiome. Lastly, we demonstrated that host specificity of *T. trichiura* is not determined by the host tissue, as first-stage (L1) larvae readily infect mouse caecaloids *in vitro.* Collectively, our results indicate that the host specificity of the human whipworm is determined by the human intestinal microbiota.

## Materials and Methods

### Mice

WT C57BL/6N and NOD.Cg-*Prkdc^scid^ Il2rg^tm1Wjl^*/SzJ (NSG) mice harbouring murine microbiota were maintained under pathogen-free conditions.

HMA mice were generated as previously described ^27^. Briefly, freshly collected faeces from healthy human donors not exposed to antibiotics for at least 6 months, hereafter referred to as ‘Donor 2’ and ‘Donor 7’, were processed within 1 h from delivery to the laboratory. The sample was homogenised at 100 mg/mL in 1x Dulbecco’s phosphate-buffered saline (D-PBS) in an anaerobic cabinet (80% CO_2_, 10% H_2_, 10% N_2_). Male and female C57BL/6N germ-free mice, bred at the Wellcome Sanger Institute, were inoculated by oral gavage with 200 µL of fresh human faecal homogenate from Donor 2 and Donor 7 once a week, for three weeks. Thereafter, these mice: ‘Donor 2 HMA’ (D2 HMA) and ‘Donor 7 HMA’ (D7 HMA), were removed from the isolator in sealed ISOcages and maintained on a positive pressure ISOrack (Tecniplast). HMA mice used in this study belonged to the fourth generation of breeding animals.

All mice were housed under a 12-h light/dark cycle at 19–24°C and 40–65% humidity. Mice were fed a regular autoclaved chow diet (LabDiet) and had *ad libitum* access to food and water. All efforts were made to minimise suffering by considerate housing and husbandry. Animal welfare was assessed routinely for all mice involved. Experiments were performed under the regulation of the UK Animals Scientific Procedures Act 1986 under the Project licence P77E8A062 and were approved by the Wellcome Sanger Institute Animal Welfare and Ethical Review Body.

### *Trichuris muris* life cycle and antigen

Maintenance of *T. muris* was conducted as described ^30^. Briefly, NSG mice (6–8 weeks old) were orally infected under anaesthesia with isoflurane with a high (n ≈ 400 eggs) dose of embryonated eggs from *T. muris* E-isolate. Mice were monitored daily for general condition and weight loss. Thirty five days later, mice were culled by cervical dislocation and the caecum and proximal colon were removed. The caecum was split and washed in RPMI-1640 plus 500 U/mL penicillin and 500 μg/mL streptomycin (all from Gibco). Worms were removed using fine forceps and cultured for 4 h or overnight in RPMI-1640 plus 500 U/mL penicillin and 500 μg/mL streptomycin at 37°C, 5% CO_2_. Media where the worms were cultured, containing eggs and excretory/secretory (ES) products, was centrifuged at 720 *g*, for 10 min, at 21–22°C (room temperature, RT), with no brake, to pellet the eggs. The eggs were allowed to embryonate for eight weeks in distilled water in the dark at RT ^11,31^. Upon completion of embryonation, eggs were long-term stored at 4°C, and infectivity was determined based on worm burdens in NSG mice infected with a high dose of eggs and culled at day 35 post infection (p.i.). The ES soluble antigen was prepared as described ^32^.

### *Trichuris trichiura* egg isolation and embryonation

*Trichuris trichiura* eggs were isolated from the faeces of an infected individual ^33^ as described below. Faeces were collected using the Fe-Col Basic Faecal collection kit (Alpha Laboratories), and 10 mL of 0.1 M sulphuric acid (Merck) was added to the collection tube to prevent bacterial and fungal growth during storage (for two weeks at RT). Faecal acid slurry was diluted in 200 mL of distilled water and passed through a series of 100 µm, 70 µm and 40 µm pluriStrainers (pluriSelect) to remove faecal debris, before being collected on a 20 µm pluriStrainer. Next, eggs were cleaned of small faecal debris using a modified “ethyl acetate flotation” protocol ^34^. Briefly, one volume of ethyl acetate (Sigma) was added to three volumes of egg suspension and thoroughly mixed. Debris was separated out by centrifugation at 3220 *g* for 5 min at RT. The aqueous phase containing debris was discarded and the egg pellet washed three times by centrifugation at 720 *g* for 10 min at RT with no brake. Eggs were resuspended in 0.01 M sulfuric acid and embryonated at 25–28°C in the dark for 3-6 weeks. Upon completion of embryonation, eggs were long-term stored at 4°C.

### Trichuris muris and T. trichiura infection of mice

Female and male WT, D2 and D7 HMA mice aged between 6 and 20 weeks old were orally infected under anaesthesia with isoflurane with a low dose (n ≈ 20) of either *T. muris* or *T. trichiura* eggs. Mice were monitored daily for general condition and weight loss. At days 35 p.i., blood was obtained by cardiac puncture under terminal anaesthesia, mice were cervically dislocated, and the mesenteric lymph nodes and caecum were collected. Serum was separated from the blood by centrifugation at 14,000 *g* for 10 min at RT and stored at −20**°**C. The caecum was slit longitudinally, and adult worms were individually removed from the caecal tissue and counted.

### Detection of parasite specific antibodies by ELISA

Levels of parasite-specific immunoglobulin IgG1 and IgG2a/c were determined by ELISA in the serum of infected WT and HMA mice as described before ^31^. Briefly, ELISA plates (Nunc Maxisorp, Thermo Scientific) were coated with 5 μg/mL *T. muris* overnight-ES soluble antigen. Serum was diluted from 1/20 to 1/2560, and parasite-specific IgG1 and IgG2a/c were detected with biotinylated anti-mouse IgG1 (Biorad) and biotinylated anti-mouse IgG2a/c (BD PharMingen) antibodies, respectively.

### Histology

To evaluate disease pathology, caecal segments were fixed in 4% paraformaldehyde and 2-5 μm paraffin sections were stained in Periodic Acid-Schiff (PAS) and Alcian Blue according to standard protocol. Images were acquired on a 3D-Histech Pannoramic-250 microscope slide-scanner using a 10X objective (Zeiss) and analysed using the NDP View2 software. Crypt lengths were measured and goblet cells per crypt counted on ∼20 crypts per animal.

### Collection, processing and culture of whole caecum, caecal luminal contents and mucosal scrapings from naive WT and HMA mice

Whole caecum samples were collected by cutting the caecum in half longitudinally to expose the whole surface of caecal contents and tissue and placing it in a Lysing Matrix E tube (MPBio). Samples were immediately stored at −80°C until DNA extraction.

Caecal luminal contents and mucosal scrapings were prepared from naive WT and HMA mice adapting protocols previously described for pigs ^25^. Briefly, the caecum was opened with curved surgical scissors, and luminal contents were scooped out and thoroughly mixed in 5 mL RPMI-1640 containing 1% L-glutamine (Gibco). Empty caecal tissue was washed with 10 mL D-PBS (Gibco) at 37°C, and mucus was gently scraped off with a cell scraper (Costar) and thoroughly mixed in 5 mL of RPMI-1640 containing 1% L-glutamine. Aliquots (200 μL) of the caecal luminal contents and mucosal scrapings were placed in a Lysing Matrix D tube (MBio), snapped frozen in liquid nitrogen and immediately stored at −80°C until DNA extraction. Bacterial metascrapes were generated by plating thoroughly mixed luminal contents or mucosal scrapings directly onto Yeast extract-Casein hydrolysate-Fatty Acids (YCFA) agar and culturing them under standard aerobic (37°C, 5% CO_2_) and anaerobic (37°C, anaerobic gas =10% H_2_, 10% CO_2_, 80% N_2_ in a Whitley DG250 workstation) conditions for 24 h. The growth on the bacterial metascrapes was then mixed with 1 mL sterile D-PBS (Gibco), scraped off, and immediately frozen at −80°C until DNA extraction ^35^.

### Bacterial culture

Bacterial culture for aero-tolerant isolates (*E. coli, Citrobacter freundii* and *Klebsiella michiganensis*) was carried out under standard aerobic conditions in Luria Bertani (LB) medium broth and agar plates. Liquid cultures were shaken at 180 rpm overnight. Aero-sensitive bacteria species (*Erysipeloclostridium ramosum*, *Phocaeicola vulgatus* and *Bacteroides uniformis*) were cultured under anaerobic conditions in YCFA medium broth and agar plates overnight ^36,37^.

### *In vitro* hatching assays with bacteria and caecal microbiota samples

Whipworm eggs were seeded in 96-well plates (Costar) at a density of 1000–3000 eggs per mL (50–150 eggs in 50 µL) for *T. muris* and 1000 eggs per mL (50 eggs in 50 µL) for *T. trichiura*. Homogenised caecal luminal contents or mucosal scrapings or an overnight culture of *E. coli* (positive control for *T. muris* egg hatching), *C. freundii, K. michiganensis, E. ramosum*, *P. vulgatus* and *B. uniformis* were added in a volume of 150 µL to each well. Each condition was run in triplicate. Plates were incubated for 24 h under aerobic or anaerobic conditions described above. Co-cultures of *T. trichiura* eggs and aero-tolerant and aero-sensitive bacteria were further incubated for three and six days.

To investigate the effect of proteases on hatching, complete Mini EDTA-free Protease Inhibitor Cocktail (Roche) was prepared according to manufacturer’s instructions and added at 2x concentration to co-cultures of whipworm eggs with *E. coli* or homogenised caecal luminal contents or mucosal scrapings.

For all hatching experiments, results were reported as percentage hatching where: Percentage Hatching = (hatched larvae / [hatched larvae + unhatched embryonated eggs]) x 100.

### *In vitro* hatching of *T. trichiura* with bleach

*Trichuris trichiura* eggs were hatched using 32% sodium hypochlorite in distilled water for 2 h at 37°C, 5% CO_2_. Eggs were washed with RPMI-1640 containing 500 U/mL penicillin and 500 μg/mL streptomycin and 10% fetal bovine serum (FBS) (all from Gibco) and incubated at 37°C, 5% CO_2_ for four to five days until hatching was observed ^11^.

### 3D Caecaloid culture

Mouse 3D caecaloids lines from C57BL/6N adult mice (6-8 weeks old) were derived from caecal epithelial crypts as previously described ^38^. Briefly, the caecum was cut open longitudinally and luminal contents removed. Tissue was cut in 0.5–1 cm segments that were washed with ice-cold D-PBS and vigorous shaking to remove mucus, and treated with Gentle Cell Dissociation Reagent (STEMCELL Tech) for 15 min at RT with continuous rocking. Released crypts were collected by centrifugation, washed with ice-cold D-PBS, resuspended in 200 μL of cold Matrigel (Corning), plated in 6-well tissue culture plates and overlaid with a Wnt-rich medium containing base growth medium (Advanced DMEM/F12 with 2 mM Glutamine, 10 mM HEPES, 100 U/mL penicillin, 100 μg/mL streptomycin, 1X B27 supplement, 1X N2 supplement (all from Gibco Thermo Fisher Scientific)), 50% Wnt3a-conditioned medium (Wnt3a cell line, kindly provided by the Clevers laboratory, Utrecht University, Netherlands), 10% R-spondin1 conditioned medium (293T-HA-Rspo1-Fc cell line, Trevigen), 1 mM N-acetylcysteine (Sigma-Aldrich), 50 ng/mL rmEGF (Gibco Thermo Fisher Scientific), 100 ng/mL rmNoggin (Peprotech), 100 ng/mL rhFGF-10 (Peprotech) and 10 μM Rho kinase (ROCK) inhibitor (Y-27632) dihydrochloride monohydrate (Sigma-Aldrich). Caecaloids were cultured at 37°C, 5% CO_2_. The medium was changed every two days and after one week, Wnt3a-conditioned medium was reduced to 30% and penicillin/streptomycin was removed (expansion medium). Expanding caecaloids were passaged, after recovering from Matrigel using Cell Recovery Solution (Corning), by physical dissociation through vigorous pipetting with a p200 pipette every six to seven days.

### Caecaloid culture in 2D conformation using transwells

3D caecaloids grown in expansion medium for four to five days after passaging were dissociated into single cells by TrypLE Express (Gibco Thermo Fisher Scientific) digestion. Two-hundred thousand cells in 200 µL base growth medium were seeded onto 12 mm transwells with polycarbonate porous membranes of 0.4 µm (Corning) pre-coated with 50 mg/mL rat tail collagen I (Gibco Thermo Fisher Scientific). Cells were cultured with expansion medium in the basolateral compartment for two days. Then, basolateral medium was replaced with medium containing 10% Wnt3a-conditioned medium for an additional 48 h. To induce differentiation of cultures, medium in the apical compartment was replaced with 50 μL base growth medium, and medium in the basolateral compartment containing 2.5% Wnt3A-conditioned medium that was changed every two days. Cultures were completely differentiated when cells pumped the media from the apical compartment and cultures looked dry.

### *Trichuris muris* L1 larvae infection of caecaloids grown in transwells

Differentiated caecaloid cultures in transwells were infected with 50–150 L1 *T. trichiura* larvae obtained by *in vitro* hatching eggs in the presence of bleach. Larvae in a volume of 100 μL of base growth medium were added to the apical compartment of the transwells. Infections were maintained for 72 h at 37°C, 5% CO_2_.

### Immunofluorescence (IF) staining of caecaloids

For IF, caecaloid cultures in transwells were fixed with 4% Formaldehyde, Methanol-free (Thermo Fisher) in D-PBS for 20 min at 4°C, washed three times with D-PBS and permeabilised with 2% Triton X-100 (Sigma-Aldrich) 5% FBS in D-PBS for 1 h at RT. Caecaloids were then incubated with primary antibodies α-villin (1:100, Abcam, ab130751), α-Ki-67 (1:250, Abcam, ab16667), α-chromogranin A (1:50, Abcam, ab15160), α-Dcamlk-1 (1:200, Abcam, ab31704), and the lectins *Ulex europaeus* agglutinin - Atto488 conjugated (UEA, 1:100, Sigma-Aldrich, 19337) and *Sambucus nigra -* Fluorescein conjugated (SNA, 1:50, Vector Laboratories, FL-1301) diluted in 0.25% Triton X-100 5% FBS in D-PBS overnight at 4°C. After three washes with D-PBS, caecaloids were stained with secondary antibody Donkey anti-rabbit IgG Alexa Fluor 555 (1:400, Molecular Probes, A31572), phalloidin Alexa Fluor 647 (1:1000, Life Technologies, A22287) and 4’,6’-diamidino-2-phenylindole (DAPI, 1:1000, AppliChem, A1001.0010) at RT for 1 h. Transwell membranes were washed three times with D-PBS and mounted on slides using ProLong Gold anti-fade reagent (Life Technologies Thermo Fisher Scientific). Confocal microscopy images were taken with a Leica SP8 confocal microscope and processed using the Leica Application Suite X software.

### DNA extraction, metagenomic library preparation and sequencing

DNA was extracted from the caecal samples (whole, mucosal scraping and luminal content) described above using FastDNA Spin Kit for Soil (MPBio) according to manufacturer’s instructions. DNA was stored at −20°C until metagenomic library preparation.

Metagenomic libraries were prepared using the Ultra II DNA Library preparation kit (NEB) and TruSeq TSQ adapters (Integrated DNA technologies). DNA concentrations were measured using the Agilent D5000 ScreenTape System (Agilent) and samples were sequenced using paired-end (2 x 150 bp) metagenomics sequencing on a HiSeq 4000 platform (Illumina).

### Metagenomic analysis

#### Metagenomic data processing

Raw sequence reads were quality-filtered using TrimGalore v0.6.0 ^39^. Contaminating mouse reads were removed by aligning reads with BWA MEM v0.7.16a-r1181 ^40^ against the mouse reference genome GRCm39.

#### Taxonomic annotation of the bacteriome

Quality-controlled sequencing reads from bulk metagenomes were taxonomically annotated at the species level by MetaPhlAn 4 v4.0.240 ^41^ using the default settings. Next, the MetaPhlAn output was converted to a taxonomy profile based on the Genome Taxonomy Database (GTDB) with GTDB-Tk v2.3.2 ^42^.

#### Microbiome community differences

Diversity estimates were inferred from the species relative abundance data generated by MetaPhlAn using the Vegan ^43^ R package. Alpha diversity was calculated using the Shannon index. Beta diversity between samples was estimated using the Bray-Curtis distance. Using a linear mixed effect model, implemented in the MaAsLin2 package in R v1.14.1^44^, differentially abundant species were identified from the WT, D2 and D7 HMA caecal microbiomes, as well as their luminal contents, and their mucosal scrapings before and after aerobic or anaerobic culture. Each of these variables (‘experimental group’, ‘sample source’ and ‘culture conditions’) were set as fixed effects in their respective analysis with MaAsLin2. A minimum prevalence threshold of 1% was set for bacterial feature selection from metagenomes, while relative abundances were log transformed without normalisation. Multiple testing correction with Benjamini-Hochberg was performed to control the false discovery rate (FDR) and the resulting associations were considered significant if *p* < 0.05.

### Statistical analysis

The Prism 10 software (GraphPad) was used to make statistical comparisons of egg hatch data. Comparisons between two groups (i.e., the absence and presence of protease inhibitor) were performed using Mann–Whitney U two-tailed tests. Kruskal Wallis and Dunn’s comparison tests were used for comparisons between three or more groups. Metagenomic sequencing data was analysed in R (v.4.4.1) as described above. Kruskal Wallis and Dunn’s comparison tests were applied to compare alpha diversity between groups.

## Results

### Colonisation of germ-free mice with human microbiota promotes *T. trichiura* infection

Studies on *T. muris* ^11,19,45–47^, *T. suis* ^24,25^ and *T. trichiura* ^47^ suggest that specific interactions between whipworm eggs and host intestinal microbiota trigger hatching and thereby determine infection establishment and host specificity in *Trichuris* species. Thus, we hypothesised that colonisation of the mouse GIT with human intestinal microbiota would enable the establishment of *T. trichiura* infection in mice. To test this hypothesis, we infected two lines of HMA mice (D2 and D7), along with WT mice harbouring a murine intestinal microbiome, with a low dose of *T. muris* or *T. trichiura* eggs *(Figure 1)*. On day 35 p.i., we evaluated the establishment of infection by measuring worm burdens and parasite-specific antibody titers (IgG1 and IgG2a/c) in the serum.

**Figure 1.**
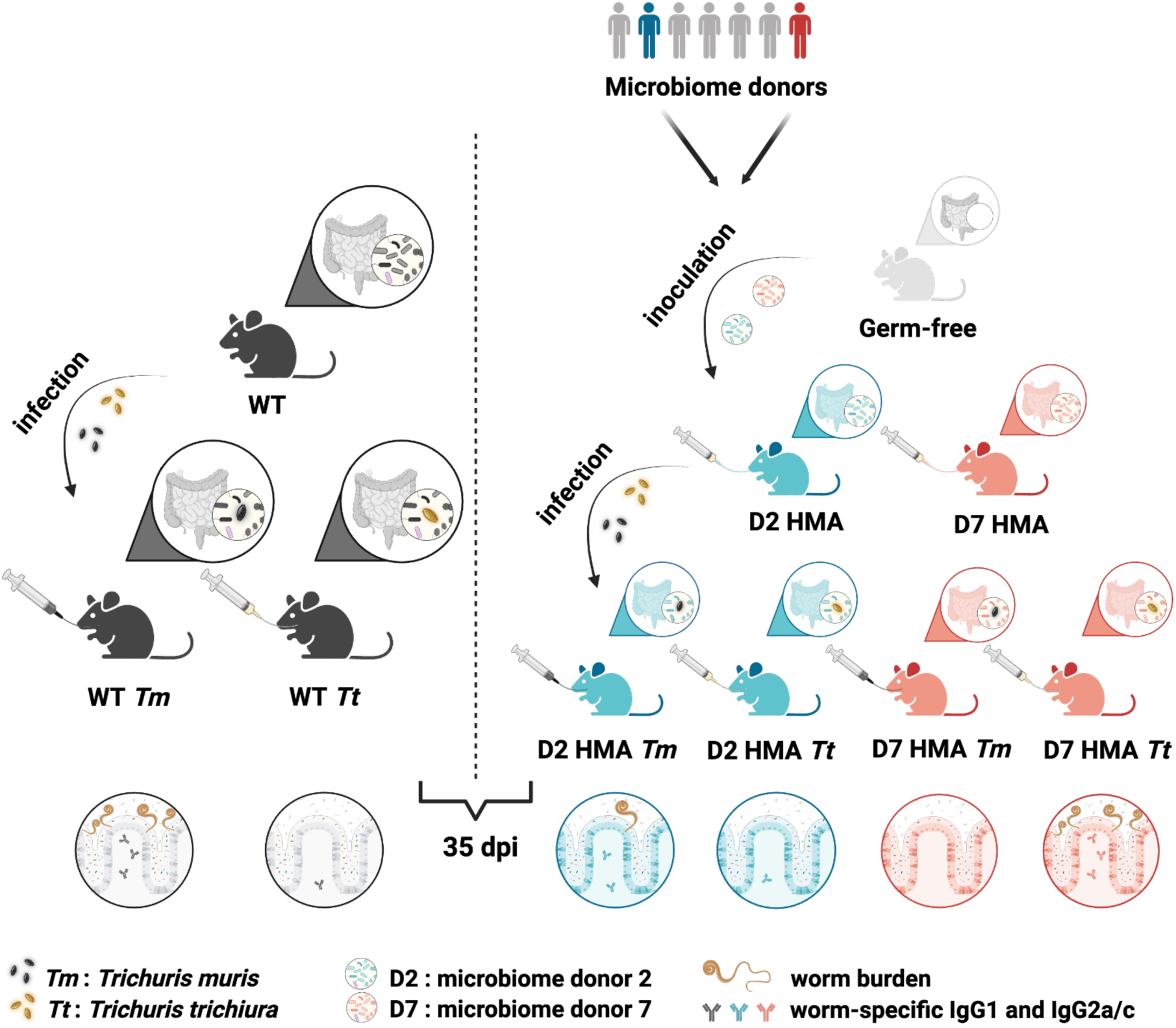
Experimental design for *Trichuris muris* and *Trichuris trichiura* infections of wild-type and human microbiota-associated mice. Experimental groups of this study consisted of C57BL/6 mice harbouring murine microbiome (wild-type, WT) and two distinct human microbiota-associated (HMA) mouse lines, each generated by inoculating C57BL/6 germ-free mice with the microbiome of a different healthy human donor (‘Donor 2’ = D2 and ‘Donor 7’ = D7). To evaluate the effect of the host microbiome on the establishment of whipworm infections, WT, D2 HMA and D7 HMA animals were orally infected with a low dose (n ≈ 20) of *T. muris* or *T. trichiura* eggs. After 35 days of infection, worm burdens (shown as worm icons) and parasite-specific antibodies (shown as Y-shaped icons) were measured. Created with BioRender.com

As expected ^48^, *T. muris* established a chronic infection in WT mice, accompanied by a mixed type 1 and 2 immune response, evidenced by the presence of both parasite-specific IgG1 and IgG2a/c in their serum (*Figure 2A–F*). In contrast, *T. muris* adult worms were only found in two out of six D2 HMA mice (*Figure 2A*) and one out of six D7 HMA mice (*Figure 2D*). No *T. trichiura* worms were recovered from either WT or D2 HMA animals (*Figure 2A*). However, we found parasite-specific antibodies in half of *T. trichiura* infected D2 HMA mice *(Figure 2B–C)*. Strikingly, *T. trichiura* infections were sustained in D7 HMA mice, with all mice harbouring worms and showing parasite-specific antibodies in their serum, at higher levels than seen for D2 HMA mice (*Figure 2D–F*). Reflecting the presence of worms in their caecum, WT mice infected with *T. muris*, and D7 HMA mice infected with *T. trichiura*, displayed longer crypts and increased numbers of goblet cells than D7 HMA mice infected with *T. muris (Figure 2G–K).* Our results, therefore, indicate that the host intestinal microbiome determines the successful establishment of whipworms in their host, with murine microbiota supporting the infection by *T. muris*, and human microbiota being required for *T. trichiura* colonisation of the mouse caecal epithelia.

**Figure 2.**
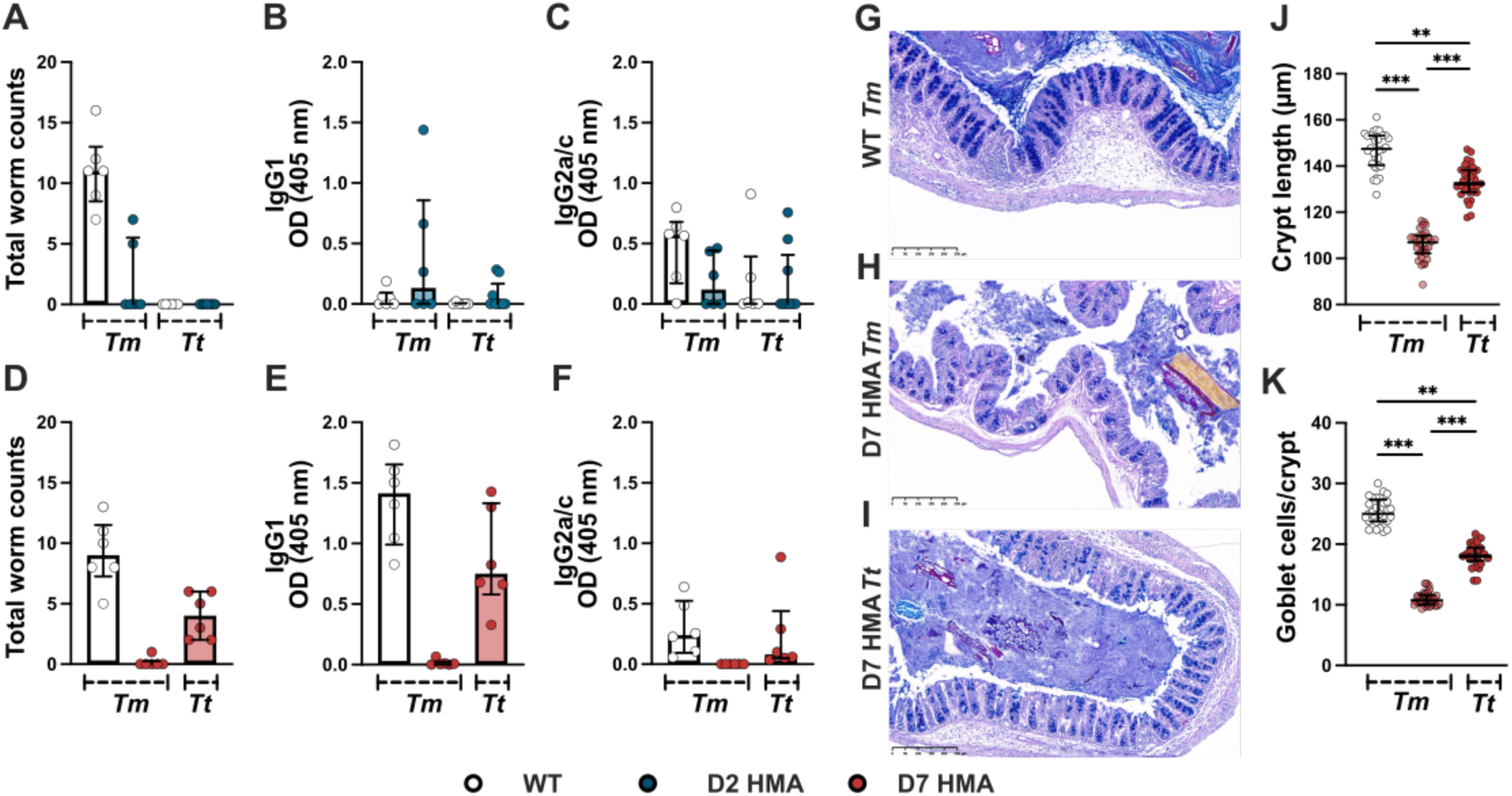
Colonisation of germ-free mice with human microbiota supports infection with Trichuris trichiura. (A–F) Worm burdens and parasite-specific antibody (IgG1 and IgG2a/c) levels in 1:40 diluted serum of *Trichuris muris (Tm)* or *T. trichiura (Tt)*-infected 5–16-wk-old wild-type (WT) and Donor 2 (D2) (A–C) or Donor 7 (D7) (D–F) human microbiota-associated (HMA) mice after 35 days of low dose infection (n ≈ 20 eggs). (G–K) Caecal histopathology of *T*. *muris*-infected WT and D7 HMA, and *T. trichiura*-infected D7 HMA mice at day 35 post infection. (G–I) Representative images from sections stained with Periodic Acid-Schiff and Alcian Blue. Scale bars 250 μm. (J) Crypt length and (K) goblet cell numbers per crypt were blindly calculated for each section (n = 5–6 mice per group, ∼ 20 crypts per mouse). Data from two independent experiments. In (A–C) WT infected with *T. muris* or *T. trichiura* and D2 HMA infected with *T. muris* n = 3 males, 3 females, D2 HMA infected with *T. trichiura* n = 4 males, 5 females. In (D–F), (J) and (K) n = 6 males for each group. Median and interquartile ranges are shown, and statistical differences between groups were evaluated using Kruskal Wallis with Dunn’s multiple comparison tests performed (\*\**p*<0.001, \*\*\**p*<0.0001).

### Successful *T. trichiura* infection is associated with the presence of unique bacterial species in the caecal microbiome of HMA mice

Our results suggest that specific members of the mouse or human gut microbiome promote the successful establishment of *T. muris* and *T. trichiura* infections in mice, respectively. Thus, to identify bacterial species potentially responsible for the differential susceptibility of mice to either mouse or human whipworms, we next characterised the caecal microbiome of WT and HMA mice using shotgun metagenomic sequencing.

Principal Coordinates Analyses (PCoA) of the caecal microbial profiles of WT, D2 and D7 HMA mice revealed samples clustered according to their host microbiome (mouse *vs* human) (PC1) and the donor (PC2), which indicate differences in bacterial species composition of the microbiota of the different mouse lines (*Figure 3A*). The alpha diversity (species-diversity within each group, measured using the Shannon-index that accounts for both the number of different species and their relative abundance) was lower in the caecal microbiome of both HMA mouse lines than the WT mice (*Figure 3B, Supplementary Table 1*), which was reflected in the total number of bacterial species comprising the murine (mean ± standard deviation (SD): 219 ± 14.85) and the humanised (D2 HMA: 79.33 ± 6.74, D7 HMA: 70.25 ± 8.04) microbiomes. This decrease in species diversity is expected as engraftment of bacterial populations in the gut of germ-free mice is impacted by exposure of aero-sensitive members of the microbiome to oxygen, low pH in the stomach, murine diet, and the genetics of the recipient mouse, among other factors ^49^. Nevertheless, the microbiomes of D2 and D7 HMA mice were distinct, composed of one-third to a half of species present in WT mice, sharing less than half of species from human microbiota origin, and containing 27 and 50 unique species, respectively (*Figure 3C, Supplementary Table 2*). Species unique to the D7 HMA microbiota, ranked by statistical significance based on a differential abundance analysis with MaAsLin2, include COE1 sp910579275 (with over 5% mean abundance)*, Bacteroides cellulosilyticus, Bacteroides clarus* and *Bacteroides xylanisolvens* (*Figures 3D-G, Supplementary Table 1* and *3)*. Collectively, these results show that D7 HMA mice host a unique caecal microbiome that may determine the establishment of *T. trichiura* infection in mice, potentially by providing critical cues for egg hatching and parasite development in the host.

**Figure 3.**
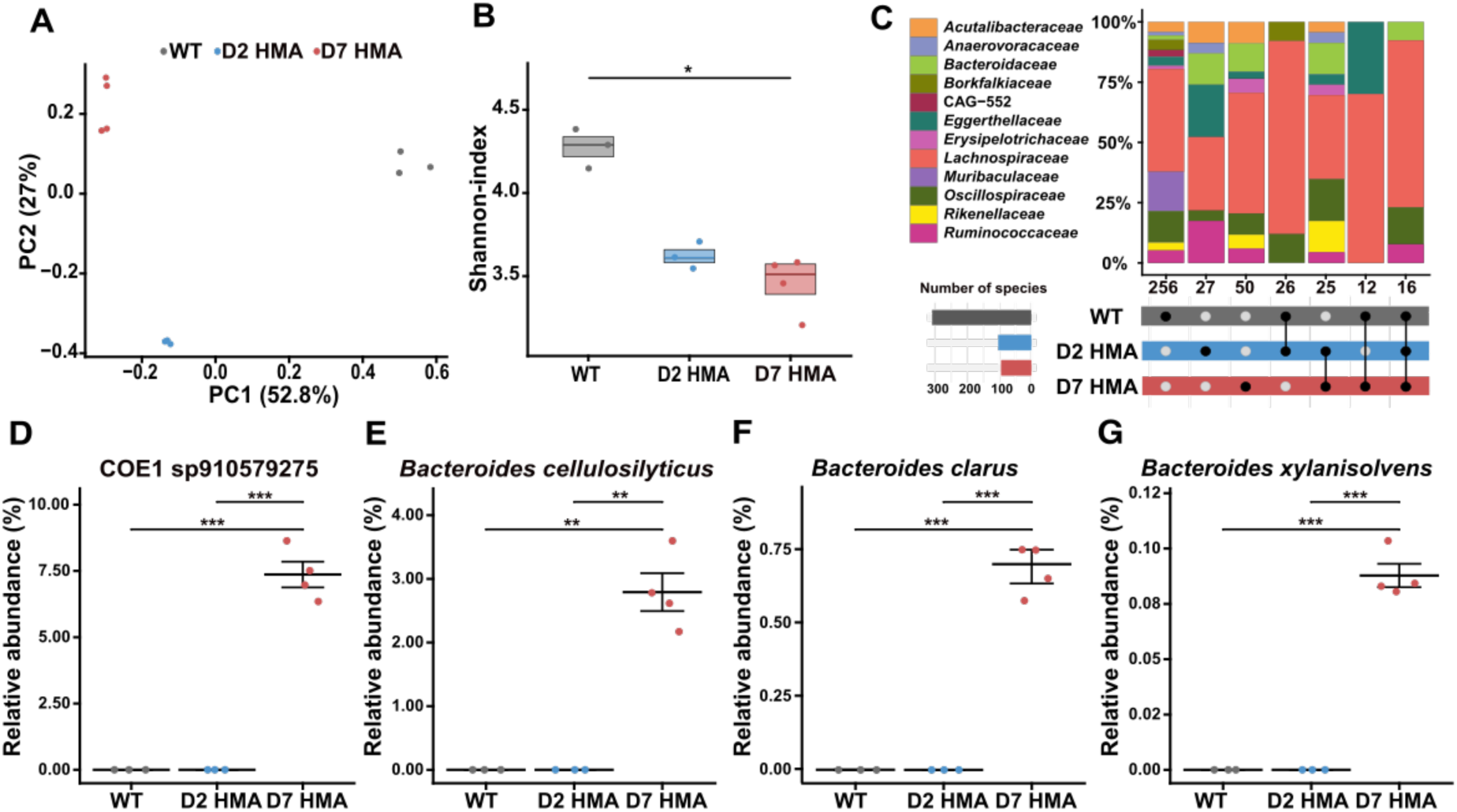
Establishment of *Trichuris trichiura* infection is associated with the presence of unique bacterial species in the microbiome of human microbiota-associated mice. Caecal microbial community structure at species level of uninfected wild-type (WT) and Donor 2 (D2) and Donor 7 (D7) human microbiota-associated (HMA) mice (WT and D2 HMA mice n = 3 per group, D7 HMA mice n = 4). (A) Principal Coordinates Analyses (PCoA) of microbial community profiles. Numbers in brackets indicate the percentage variance explained by that component. (B) Alpha-diversity (Shannon) index at species level. Median and interquartile range are shown. Statistical differences between groups were evaluated using Kruskal Wallis test and pairwise comparisons were conducted using Dunn’s test (\**p*<0.05). (C) Upset plot showing unique and shared microbiome species between mouse lines. Vertical bar plots represent the percentages of species, coloured by family, present in the mouse lines highlighted in the lower panel. Numbers below the bars indicate the number of species in each intersection. Horizontal bar plots on the left side panel show the total number of species detected in each mouse line. (D–G) Relative abundance of COE1 sp910579275*, Bacteroides cellulosilyticus, Bacteroides clarus* and *Bacteroides xylanisolvens* in the caecal microbiome of WT, D2 HMA and D7 HMA mice. These bacteria are significantly enriched in D7 HMA mice compared to WT and D2 HMA mice as determined by Multivariable Association with Linear Models (MaAsLin). Mean and standard error of mean are shown (\*\**p*<0.001, \*\*\**p*<0.0001).

### Microbial populations within the caecal mucosa from HMA mice induce *in vitro* hatching of *T. trichiura* eggs

The presence of unique bacterial species in the microbiome of WT and HMA mice led us to hypothesise that successful colonisation and specificity of *T. muris* for WT mice versus that of *T. trichiura* for D7 HMA mice is partly a consequence of hatching of the eggs in response to the intestinal microbiota from their specific hosts. Importantly, the outer mucus layer of the large intestine hosts a microbiota distinct from that of the lumen ^50,51^, and hatching of *T. suis* is induced by the mucosal scrapings from the pig caecum and colon ^25^. Therefore, to test our hypothesis, we evaluated *in vitro* hatching of *T. muris* and *T. trichiura* eggs in the presence of mucosal scrapings and luminal contents from the caecum of naive WT and HMA mice under aerobic and anaerobic conditions (*Figure 4*). Co-culture with *E. coli* served as a positive control for *T. muris* hatching (*Supplementary Figure 1*).

**Figure 4.**
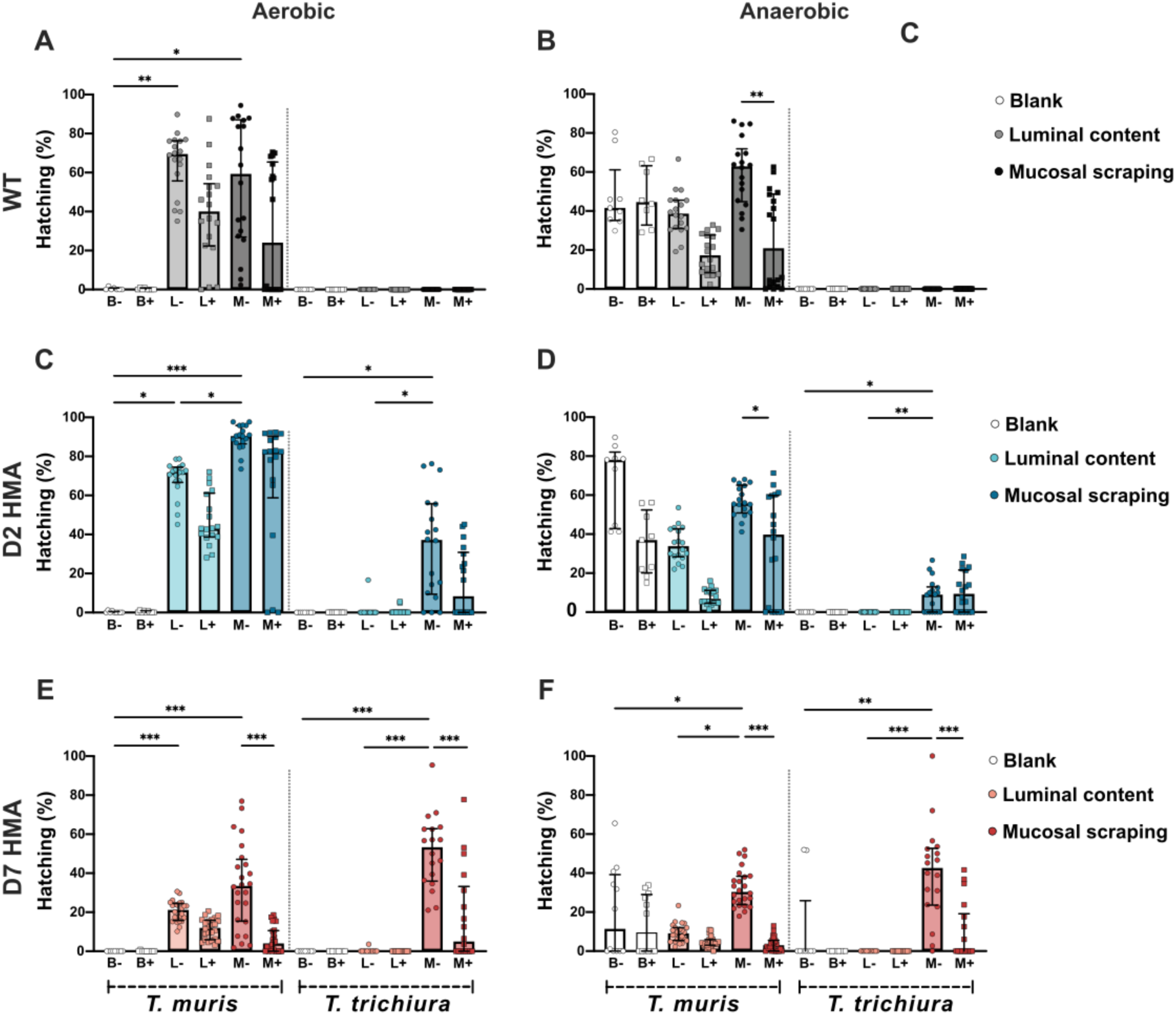
The microbiome of human microbiota-associated mice (HMA) triggers *in vitro* hatching of *Trichuris trichiura* eggs through protease-mediated mechanisms. *Trichuris muris* or *T. trichiura* eggs were co-cultured with RPMI-1640 media (blank, B), caecal luminal contents (L) or mucosal scrapings (M) from (A and B) wild-type (WT), (C and D) Donor 2 (D2) and (E and F) Donor 7 (D7) HMA mice in the absence (-) or presence (+) of a protease inhibitor cocktail (2x) under aerobic or anaerobic conditions at 37°C for 24 h. The number of total embryonated eggs and hatched larvae were counted, from which the percentage of hatching was calculated. Hatching was completed in triplicate across 2-4 independent experiments, with each independent experiment involving 2 animals as sample sources, and each dot representing a well of eggs (n = 12-24). Median and interquartile range are shown, and statistical differences between groups were evaluated using Kruskal Wallis with Dunn’s multiple comparison tests (\**p*<0.05, \*\**p*<0.001, \*\*\**p*<0.0001).

*T. muris* eggs hatched in response to mucosal scrapings and luminal contents of the caecum of both WT and HMA mice (*Figure 4*). Surprisingly, the highest levels of *T. muris* hatching were observed in response to samples from D2 HMA mice (*Figures 4C and D*), followed by co-culture with *E. coli* (*Supplementary Figure 1*) and mucosal scrapings and luminal contents of WT mice (*Figures 4A and B*). The lowest levels of *T. muris* hatching were induced by samples from D7 HMA animals (*Figures 4E and F*). These results suggest that while the caecal microbiome of D2 HMA mice harbours bacterial species capable of hatching *T. muris* eggs, this microbial community does not support the establishment and persistence of *T. muris* chronic infections (*Figures 2A-C*).

In contrast, *T. trichiura* eggs only hatched when co-cultured with mucosal scrapings from the caecum of HMA mice *(Figures 4C-F)*. No hatching was observed in response to samples from WT mice *(Figures 4A and B)* or luminal contents from HMA mice *(Figures 4C-F)*. In line with our *in vivo* infection results *(Figure 2)*, caecal samples from D7 HMA mice induced higher levels of hatching than those of D2 HMA mice *(Figures 4C-F)*, suggesting that the caecal microbiome of D7 mice contains bacterial species critical for *T. trichiura* hatching and colonisation of its host.

Hatching of both *T. muris* and *T. trichiura* was consistently higher in response to mucosal scrapings, compared to luminal contents from the same mice, and under aerobic conditions *(Figure 4).* Interestingly, we observed “spontaneous” hatching of *T. muris*, but not *T. trichiura,* in control (blank) wells where eggs were co-cultured with media under anaerobic conditions; however, no visible contamination was observed in these co-cultures (*Figures 4B, D and F*). Similar observations on hatching of *T. muris* in media have been reported by Sargsian and colleagues ^47^.

Recently, we found that hatching of *T. muris* in response to monocultures of fimbriated and non-fimbriated bacteria is protease-dependent ^23^; thus, to determine if microbiota-induced hatching of *T. muris* and *T. trichiura* is also mediated by proteolytic enzymes, we used a protease inhibitor cocktail in our *in vitro* hatching assays. We found that inhibition of proteases reduces *T. muris* and *T. trichiura* hatching in response to samples from WT and HMA mice *(Figure 4)*. However, the level of reduction is variable; hatching induced by *E. coli* (*Supplementary Figure 1*) and mucosal scrapings of D7 HMA mice *(Figures 4E and F)* was almost completely ablated by the protease inhibitor cocktail, whereas co-culture with samples from WT and D2 HMA mice only showed a slight reduction in hatching *(Figures 4A-D)*. These results likely reflect variations in the bacterial load of the GIT slurry and a dose-dependent effect of the protease inhibitors ^23^.

Taken together, these findings indicate that whipworm hatching *in vivo* is triggered by bacterial communities closely associated with the caecal epithelia, and that is a process dependent on proteases. Moreover, these findings suggest that host specificity of *Trichuris* species is a consequence of the host intestinal microbiome, which is required for the induction of egg hatching as well as for supporting the development of the parasite inside their host.

### Identification of bacterial species within the caecal mucosal niche of HMA mice that trigger hatching and determine host specificity of *T. trichiura*

To characterise the composition of the microbiota within the mucosal niches of the caecum of D7 and D2 HMA mice and identify key bacterial species responsible for *T. trichiura* hatching, we performed metagenomic sequencing on caecal mucosal scrapings and luminal contents of naive animals.

The microbiome associated with the caecal mucosa showed lower alpha diversity than that of the lumen in both HMA mouse lines *(Figure 5A, Supplementary Figure 2A)*, mirroring a decrease in the number of species present in this niche (D7 HMA luminal content, mean ± SD: 96.8 ± 1.71, D7 HMA mucosal scraping: 42 ± 6.44; D2 HMA luminal content: 85.08 ± 3.79, D2 HMA mucosal scraping: 56.5 ± 3.86) (*Supplementary Table 4 and 5)*. Notably, the mucosal scraping was composed of a subset of the species present in the luminal content (*Supplementary Figure 3*); nonetheless, 75 and 61 species showed significant differences in relative abundance in the luminal versus the mucosal microbiome of the caecum of D7 and D2 HMA mice, respectively (*Supplementary Tables 4-7*). The luminal contents of both D2 and D7 HMA mice showed high abundance of *B. uniformis* and *Eubacterium_F sp910575665,* but each microbiome contained also other species specifically associated with each mouse strain (*Figure 5B, Supplementary Figure 2B* and *Supplementary Tables 4-7*). In contrast, the mucosal microbiome of D2 and D7 HMA mice showed a significantly higher abundance of species/taxa from the Lachnospiraceae and Ruminococcaceae families, including UBA3282 sp910584965 and *Anaerotruncus s*p000403395, present in both mouse strains; COE1 sp000403215 and CAG-95 sp009917455, increased in D2 HMA mice; and COE1 sp910579275 and *Acetatifactor* sp910579755, increased in D7 HMA mice (*Figures 5B-F, Supplementary Figure 2B* and *Supplementary Tables 6-7*). Interestingly, COE1 sp910579275 is one of the unique species present in the microbiome of D7 HMA mice *(Figure 3D)*, strongly suggesting this bacterium contributes to the hatching and host specificity of *T. trichiura*. COE1 sp910579275 represents an uncharacterised genus of bacteria with no cultured representative species thus far. Therefore, future efforts should focus on defining the necessary culture conditions of this potentially important taxon to enable experimental studies.

**Figure 5.**
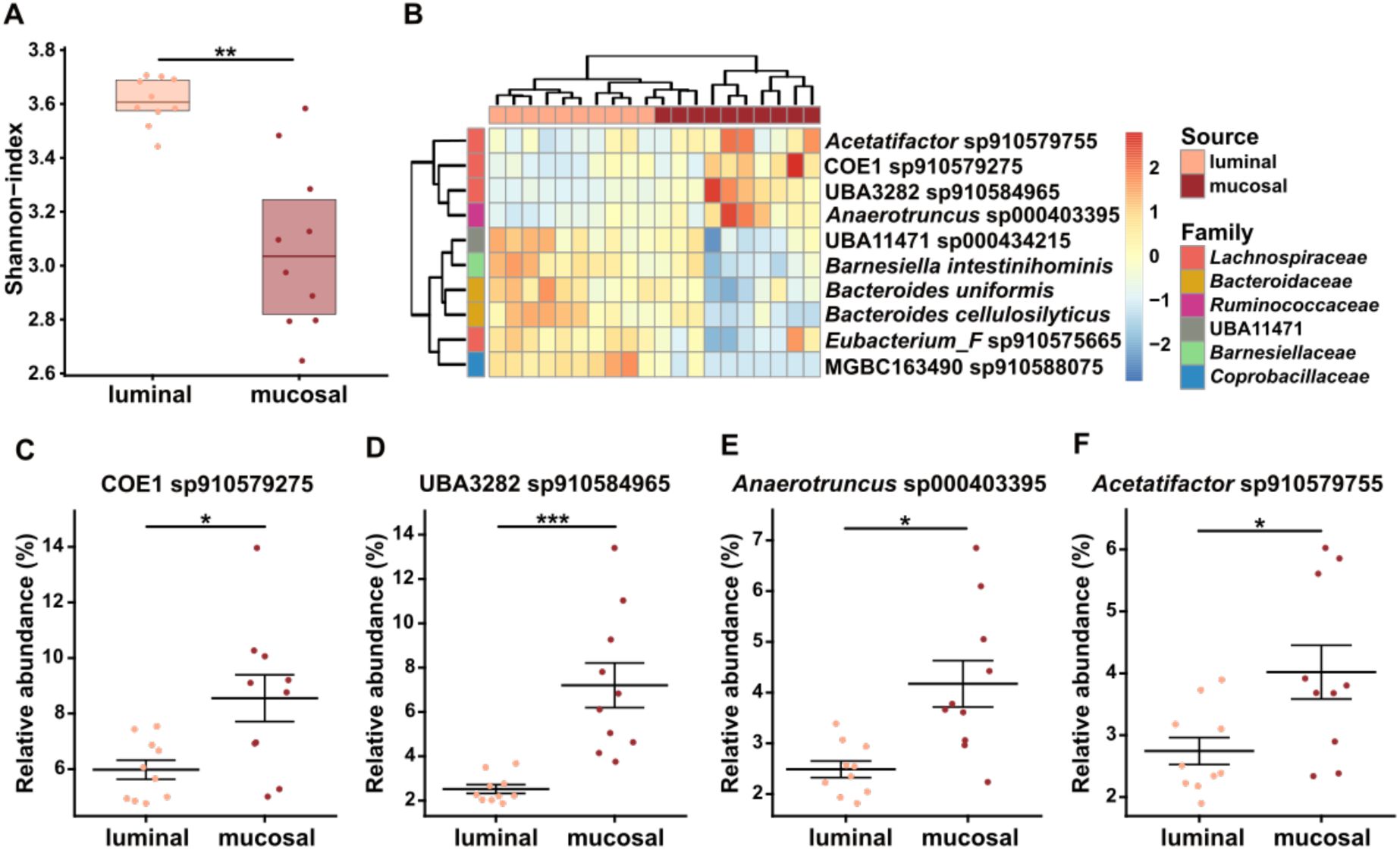
Identification of bacterial species with increased abundance within the mucosal niche of the caecum of human microbiota-associated mice, which are associated with *Trichuris trichiura* infection. Caecal microbial community structure at species level of the luminal content (luminal), and mucosal scraping (mucosal) of Donor 7 (D7) human microbiota-associated (HMA) mice (n = 10). (A) Alpha-diversity (Shannon) index. Median and interquartile range are shown. Statistical differences between groups were evaluated using Kruskal Wallis with Dunn’s multiple comparison tests (\*\**p*<0.001). (B) Heatmap depicting the relative abundance of the ten most abundant microbial species that show statistically significant differences between the lumen (luminal content) and mucosa (mucosal scraping) of the caecum in D7 HMA mice. The heatmap uses a colour scale to represent relative abundances, highlighting variations across the two microbial niches. Clustering was performed using hierarchical clustering with complete linkage on Euclidean dissimilarity matrices. Each column represents a microbial sample. (C-F) Relative abundance of COE1 sp910579275, UBA3282 sp910584965, *Anaerotruncus* sp000403395 and *Acetatifactor* sp910579755. These bacteria are significantly enriched in the caecal mucosa, compared to the lumen, of D7 HMA mice as determined by Multivariable Association with Linear Models (MaAsLin). Mean and standard error of mean are shown (\**p*<0.05, \*\*\**p*<0.0001).

Hatching of *T. trichiura in vitro* could be triggered by either: 1) bacterial species of high abundance in the uncultured initial mucosal scrape the eggs encounter at the start of the co-culture *(Figure 5B-F* and *Supplementary Figure 2B*) or 2) bacterial species that become more abundant, through growth, during the 24 hours incubation with the eggs. To identify bacterial species selectively expanded/enriched by aerobic or anaerobic culture that could induce *T. trichiura* hatching, we grew metascrapes of mucosal scrapings and luminal contents on YCFA agar and analysed their microbial composition using metagenomic sequencing. Culture of mucosal scrapings impacted the structure of their microbial communities, resulting in a significant decrease in species diversity, which was more pronounced under aerobic conditions (*Figure 6A* and *B, Supplementary Table 8, Supplementary Figure 2C* and *Supplementary Table 9).* This reduction is expected, as aerobic and anaerobic conditions selectively favour the growth of bacteria depending on their oxygen tolerance, with the majority of species of the intestinal microbiome being obligate anaerobes ^52^. Among the unculturable species, lost upon culture, are several of the highly abundant taxa of the microbiome of the mucosal scrapings of D7 HMA mice, including COE1 sp910579275, *Acetatifactor* sp910579755 and *Anaerotruncus sp000403395,* which failed to grow in both the D7 and D2 HMA mucosal metascrapes *(Supplementary Figures 4A and B)*.

**Figure 6.**
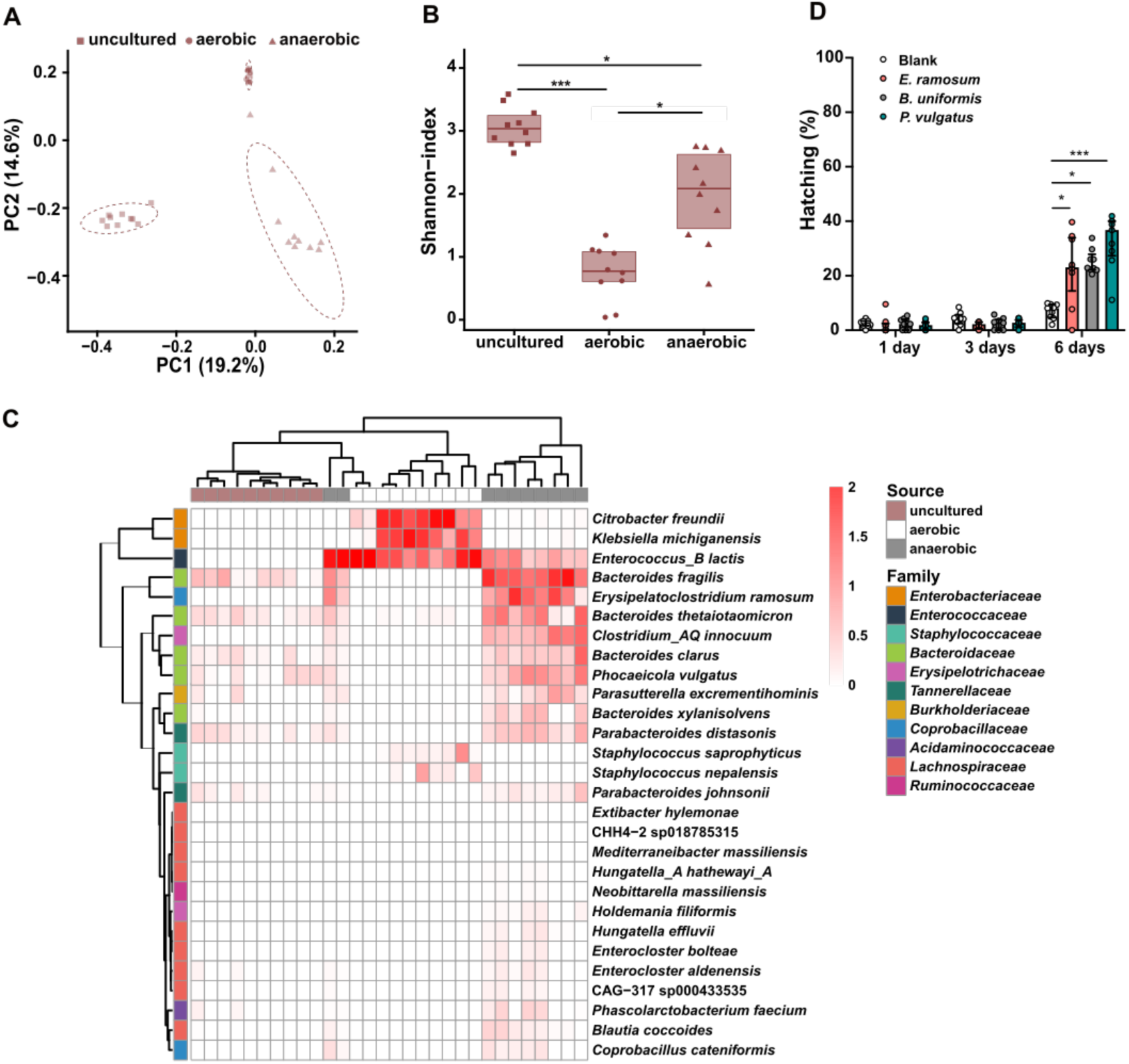
Anaerobic culture of caecal mucosal scrapings of human microbiota-associated mice increases the abundance of bacterial species that trigger *Trichuris trichiura* egg hatching. (A-C) Microbial community structure at species level of the caecal mucosa of Donor 7 (D7) human microbiota-associated (HMA) mice before (uncultured) and after 24 h of culture under aerobic or anaerobic conditions (n = 10 mice). (A) Principal Coordinates Analyses (PCoA) of microbial community profiles. Numbers in brackets indicate the percentage variance explained by that component. (B) Alpha-diversity (Shannon) index. Median and interquartile range are shown, and statistical differences between groups were evaluated using Kruskal Wallis test. Pairwise comparisons were made using Dunn’s test (\**p*<0.05, \*\*\**p*<0.0001). (C) Heatmap depicting the relative abundance of all the microbial species statistically significantly more abundant in the aerobic and anaerobic metascrapes compared to the uncultured caecal mucosal scrapings of D7 HMA mice. The heatmap uses a colour scale to represent relative abundances, highlighting variations of the microbial composition across the two culture conditions. Clustering was performed using hierarchical clustering with complete linkage on Euclidean dissimilarity matrices. Each column represents a microbial sample. (D) *Trichuris trichiura* eggs were co-cultured with Yeast extract-Casein hydrolysate-Fatty Acids media (blank), or *Erysipeloclostridium ramosum*, *Bacteroides uniformis* and *Phocaeicola vulgatus* at 37°C for one, three and six days under anaerobic conditions. The number of total embryonated eggs and hatched larvae were counted, from which the percentage of hatching was calculated. Hatching was completed in triplicate across 3 independent experiments and each dot represents a well of eggs (n = 9). Median and interquartile range are shown, and statistical differences between blank and bacteria were evaluated using Kruskal Wallis with Dunn’s multiple comparison tests performed (\**p*<0.05, \*\*\**p*<0.0001).

Aero-tolerant species that expanded under aerobic culture in the D7 HMA mucosal metascrapes include the Enterobacteriaceae *C. freundii* and *K. michiganensis* and *Enterococcus B lactis (Figure 6C, Supplementary Tables 8* and *10*). Notably, aerobic culture of the D2 HMA caecal mucosal scraping resulted in enrichment of *E. coli (Supplementary Figure 2D, Supplementary Tables 9* and *11)*, which could explain the high levels of *in vitro T. muris* egg hatching induced by microbiota from D2 HMA mice *(Figure 4C*). Anaerobic culture of HMA mucosal scrapings significantly increased the abundance of obligate anaerobes that included: several Bacteroidaceae, namely *B. xylanisolvens* and *B. clarus,* which were unique to the caecal microbiome of D7 HMA mice; *B. uniformis,* which was specifically enriched in anaerobic metascrapes from D2 HMA mice and highly abundant in D7 HMA mucosal scrapings and anaerobic cultures; *B. fragilis* and *B. thetaiotaomicron,* which were only present in the microbiota of HMA mice; and *P. vulgatus,* which showed higher abundance in HMA compared to WT mice. Further, anaerobic culture also resulted in increased abundance of a Coprobacillaceae family member, *E. ramosum,* which was solely detected in HMA mice *(Figure 6C, Supplementary Figures 2D and 5, Supplementary Tables 8-12)*.

To test if the species enriched under aerobic and anaerobic culture contribute to *T. trichiura* hatching, we performed *in vitro* hatching experiments with monocultures of bacterial species accessible to us, *C. freundii*, *K. michiganensis, P. vulgatus*, *E. ramosum* and *B. uniformis*. We and others have observed hatching of *T. muris* in as little as 40–60 min upon *in vitro* culture with mouse caecal contents ^19^, or two hours of co-culture with different fimbriated and non-fimbriated bacteria ^15–18,20–23^, and found *T. trichiura* eggs hatched after 24 h of co-culture with caecal mucosal scrapings of HMA mice *(Figure 4)*. In contrast, Sargsian and colleagues ^47^ reported hatching of *T. trichiura* after six days of incubation with monocultures of *P. sordellii*. Therefore, we evaluated levels of hatching after one, three and six days of co-culture. Aero-tolerant bacteria (*C. freundii* and *K. michiganensis)* did not trigger *T. trichiura* hatching (*Supplementary Table 13).* In contrast, co-culture for six days, but not one or three days, with the anaerobes *P. vulgatus*, *E. ramosum* and *B. uniformis* (*Figure 6D*) resulted in hatching of 37%, 23% and 23% of *T. trichiura* eggs, respectively.

Altogether, our comparative metagenomic analysis resulted in the identification of members of the caecal mucosal microbiome of HMA mice that trigger *T. trichiura* hatching, strongly suggesting they are key determinants of host specificity of *T. trichiura*.

### *Trichuris trichiura* host specificity is not determined by caecal epithelia

After hatching, whipworm L1 larvae rapidly invade host epithelial cells at the crypts of the intestinal epithelium becoming completely intracellular ^53,54^. While yet to be identified, unique host intestinal epithelial cues, such as components of the mucus layer or cell membrane receptors mediating parasite invasion of the epithelia, could contribute to *Trichuris* host specificity. To test this hypothesis, we next investigated if chemically-hatched *T. trichiura* L1 larvae could invade the mouse caecal epithelia *in vitro* using caecaloids ^53^. Strikingly, we found *T. trichiura* L1 larvae invaded mouse caecaloids forming syncytial tunnels that spanned several epithelial cells (*Figure 7, Supplementary Movie 1*), which were identical to those formed by *T. muris* L1 larvae ^53^. Together with our *in vivo* experiments (*Figure 2*), these results suggest the host intestinal epithelium cues that mediate L1 larvae penetration in the tissue are conserved among mice and human and do not play a major role on *T. trichiura* host specificity/selective infection of human and non-human primates.

**Figure 7.**
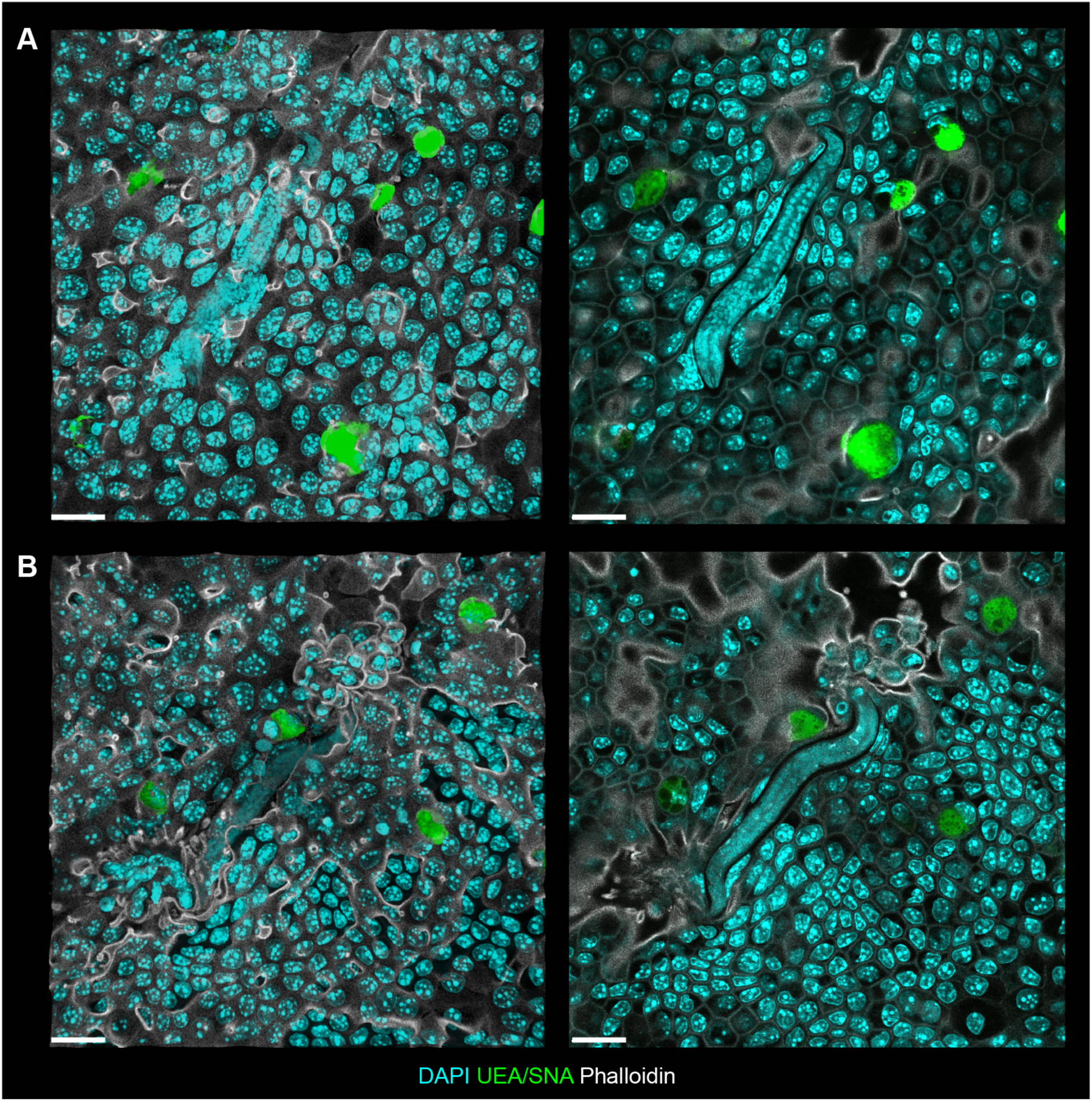
*Trichuris trichiura* L1 larvae invades mouse caecal epithelial cells. Representative confocal immunofluorescence (IF) images of mouse caecaloids infected with *Trichuris trichiura* L1 larvae for 72 h. Complete z-stack and selected individual maximum projections showing larvae infecting intestinal epithelial cells within syncytial tunnels. In green, the lectins UEA and SNA bind mucins in goblet cells; in aqua, DAPI stains nuclei of intestinal epithelial cells and larvae, respectively; and in white, phalloidin binds to F-actin. Scale bars 20 μm. IF imaging experiments on *T. trichiura-*infected caecaloids were done in triplicate across three independent replicas using two caecaloid lines derived from two C57BL/6 mice.

## Discussion

The long-term co-evolution of whipworms with their hosts has not only resulted in a marked tropism of whipworms for the caecum, where eggs passively accumulate, but also a preference for specific hosts ^12,19,55^, potentially driven by unique interactions with the diverse intestinal microbiota encountered in different mammals ^56^. Here, we have demonstrated that the host intestinal microbiome is indeed a key determinant of *T. trichiura* host specificity, by showing that colonisation of the mouse GIT with human microbiota provides the essential factors both for *T. trichiura* hatching and subsequent development inside the murine caecal epithelium. Moreover, the GIT of HMA mice does not support *T. muris* infections that normally establish in mice harbouring a murine gut microbiome. Our findings, together with previous studies on the impact of the host microbiota on parasite hatching, fitness and survival ^11,15,16,19,22,23,45–47,57^, as well as the effects of whipworm infection on the restructuring of the intestinal bacterial communities of their host ^47,58–60^, shed further light on the three-way co-adaptation processes of whipworms, their mammalian hosts and host-gut microbiomes that enable successful infections.

In our study, we took advantage of two lines of HMA mice that have been previously generated to investigate how the host microbiome influences the response to infection with another human helminth, *Schistosoma mansoni* ^27–29^. The *Schistosoma* studies revealed human gut microbiota roles in modulating susceptibility, pathology and tissue damage control during infections. Notably, these effects varied dramatically depending on the donor’s baseline microbiome composition ^27–29^. Although the availability of HMA mouse lines was limited, we were able to show that mice with microbiomes originating from two donor humans were able to support *in vitro* hatching of *T. trichiura* eggs. Furthermore, detection of parasite-specific antibodies in the serum of approximately half of the infected D2 HMA mice, and all of the infected D7 HMA mice, indicated that their microbiomes also supported the initial establishment of an infection. Consistent with the previous *Schistosoma* findings, we found that the source/donor of the human microbiome has a major impact on the host susceptibility to an infection, with D7 HMA mice consistently sustaining *T. trichiura* infections even further, so that viable worms could be recovered at day 35 p.i. Our *in vivo* infections and *in vitro* hatching assays both demonstrated that human microbiota is critical for the induction of *T. trichiura* hatching; however, they also indicated that bacterial species solely present in the caecal microbiome of the D7 HMA mice are crucial for sustaining infections beyond induction. These unique taxa included several members of the Lachnospiraceae (e.g. COE1 sp910579275) and Bacteroidaceae (e.g. *B. clarus* and *B. xylanisolvens)*, families previously associated with *T. trichiura*-colonised individuals ^47^ and with species making up the whipworm microbiome ^11^. In contrast, the microbiome of humanised-microbiota mice only sustained *T. muris* infections in very few animals, similar to observations in gnotobiotic mice containing a synthetic human gut microbiota where high-dose inoculations resulted in poor establishment by only small numbers of worms ^61^, suggesting that human intestinal microbiota members do not appropriately support *T. muris* hatching and development. The murine microbiome of WT mice, which supported *T. muris* hatching and chronic infection, presented a strikingly different microbial community structure to that of HMA mice (*Supplementary Figure 6*, Supplementary Tables 12, 14-16**) that we suggest determines host specificity for the mouse whipworm. The particular bacterial species that favour whipworms establishing infections and persisting could do so in three ways. First, by triggering egg hatching; second, by serving as a source of bacteria for the parasite’s own microbiota, which has been shown to be required for whipworm fitness ^11^; and third, by directly or indirectly (through modulation of the intestinal epithelium) providing metabolites that cannot be synthesized by the parasite but that are critical for parasite development and survival ^10^.

Given the distinctive nature of whipworm’s multi-intracellular niche in the caecal epithelium, our observations on the increased levels of *T. trichiura* and *T. muris* hatching in response to murine caecal mucosal scrapings, together with previous findings of *T. suis* hatching in response to pig intestinal mucosal scrapings ^25^, point to the evolution of mechanisms of whipworm hatching in response to bacterial communities closely associated with the intestinal epithelia. The proximity of the location of hatching to the parasite niche may facilitate rapid larval invasion of epithelial cells after eclosion and parasite survival in the oxygen depleted environment of the intestine. Interestingly, the caecal mucosal niche of HMA mice harboured a high abundance of Lachnospiraceae family members (including the already highlighted taxon COE1 sp910579275), strongly indicating these species are key triggers of *T. trichiura* hatching. Since these bacteria remain unculturable, future efforts to establish suitable culture conditions will be needed to validate our findings. Nevertheless, both aerobic and anaerobic culture of mucosal scrapings have revealed additional bacterial species that can also facilitate human whipworm hatching, including members of the Enterobacteriaceae and Bacteroidaceae families. Indeed, *P. vulgatus*, *B. uniformis* and *E. ramosum* successfully induced *T. trichiura* hatching *in vitro*, albeit after six days of co-culture. This extended co-culture that surpasses the intestinal transit time could reflect the need for a specific bacterial density threshold or the accumulation of bacterial metabolites or proteases, naturally occurring the caecum — a fermentative organ, for the activation of processes that lead to the degradation of the egg’s polar plugs ^16,23^. Of note, while in our hands *P. vulgatus* was the best hatching inducer, previous co-cultures of this bacterium with *T. trichiura* hatching did not result in hatching ^47^, potentially due to intra-strain differences. Paradoxically, while *T. trichiura* hatching in response to mucosal scraping was higher under aerobic conditions, monocultures of aerobes expanded in aerobic metascrapes did not trigger hatching *in vitro*. This finding suggests that whilst human whipworm hatching is triggered by anaerobic bacteria, it benefits from higher oxygen concentrations that can support larval activation and movement required for the degradation of the egg polar plugs and successful eclosion from the egg ^23,62^.

The mechanisms mediating whipworm larvae recognition and invasion of the intestinal epithelia, and parasite development inside this unique niche, remain unknown but could also be determinants of *Trichuris* host specificity. Thus, our results on *T. trichiura* infection of HMA mice and mouse organoids (by pre-hatched larvae) were surprising and suggest that intestinal cues/receptors, sensed by the larvae and triggering tissue penetration, are shared between mice and humans. These findings together with earlier observations by Panesar and Croll ^19^ on successful *T. muris* infections in caecatomised mice, where the parasites established chronic infections in the ileum, suggest that while intracellular infection of an “intestinal” epithelium is essential for whipworm development, the species and regional origin of the intestinal tissue do not play a role in whipworm host specificity. Instead, “the appropriate conditions for hatching, emergence” ^19^ and development of the larvae, now identified as the presence of specific bacteria of the host gut microbiome, and from which the parasite acquires their own microbiota, seem to be crucial in the establishment of patent infections with *Trichuris* species. In line with this, Beer reported *T. trichiura* L1 larvae hatching and invading the caecal epithelium of pigs, but with stunted development and ultimately being expelled ^14^, similar to our observations in D2 HMA mice where bacterial taxa critical for parasite development were missing.

To date, studies of *Trichuris*-colonised individuals from endemic regions have failed to clearly associate specific taxa in gut microbiomes with whipworm colonisation, likely due to hidden effects of diet, ethnic and environmental differences of distinct geographic regions on the microbiome of affected populations ^10^. Nevertheless, Sargasian and colleagues ^47^ recently identified OA-8, a species from the *Peptostreptococcaceae* family enriched in the faecal microbiome of helminth-infected individuals from rural Malaysia, as an inducer of *T. trichiura* and *T. muris* hatching ^47^. We did not detect any members of this family in the microbiome of HMA mice, which were generated with faeces from healthy donors living in the United Kingdom. Given the historical global prevalence of human whipworm infections ^2,4,63–68^, the microbiome-driven conditions for *T. trichiura* hatching and persistence must be ubiquitous across all human populations, instead of being limited to unique bacteria enriched in the microbiomes of communities in current disease-endemic regions. Consistent with this, whipworm infections remained frequent in developed countries, such as the UK, until the second half of the last century ^14,69^ and still can establish in individuals from these nations ^33^. Nevertheless, inter-individual microbiota differences may impact the susceptibility of individuals in a community, which could partially explain the range of infection burdens found in populations from endemic areas and our results on the differential susceptibility to *T. trichiura* between D2 and D7 HMA mice. Furthermore, our findings on the higher levels of *T. trichiura* hatching induced by bacterial communities from caecal mucosa disagree with those from metagenomic studies of faecal samples, which do not reflect the microbial populations of the human GIT, in particular those at the site of whipworm infection ^70,71^. We, however, acknowledge the limitations for the collection of human mucosal samples, which currently requires colonoscopy or alternative access to post mortem samples ^70,72^. Nevertheless, access to this material will enable investigations to further dissect the interactions between human microbiome members and *T. trichiura* and the mechanisms that mediate human whipworm hatching, development and persistence as chronic infections. Similarly, access to GIT samples of non-human primates such as baboons and macaques (from zoos or bioparks), which are also hosts of *T. trichiura*, would allow the identification of bacterial species shared between the microbiota of humans and non-human primates that determine their susceptibility to the human whipworm tying in evolutionary aspects shared by these species, their microbiome and their parasites ^68^.

In summary, we propose that host specificity of *Trichuris* species is determined by their host intestinal microbiome. A better understanding on how these co-adaptation processes impact whipworm colonisation and persistence in their hosts may reveal vulnerabilities that can be targeted for intervention of these important human and veterinary parasites. Notably, the identification of bacterial species that induce hatching of *T. trichiura* will steer future work to develop more refined gnotobiotic mouse models for *T. trichiura* infection as well as *in vitro* protocols to obtain live L1 larvae that can be used to translate our current *in vitro* caecaloid systems to the human whipworm ^38,53^. Together, these models will unlock opportunities to: 1) investigate human whipworm pathophysiology and immunity, 2) understand differences in the metabolic requirements of *T. trichiura* and *T. muris* linked to the host microbiome and potentially determining host specificity; and 3) discover and test new and effective drugs to control *Trichuris* infections.

## Acknowledgements - Funding

This work was supported by the Sir Henry Dale Fellowship jointly funded by the Wellcome Trust and the Royal Society (222546/Z/21/Z, M.A.D-C.); the Wellcome Trust (206194, M.B; 203151/Z/16/Z, 203151/A/16/Z, M.A.D-C) and the UKRI Medical Research Council (MC_PC_17230, M.A.D-C). K.A.S is supported by Cambridge Trust. The Bioimaging Facility microscopes used in this study were purchased with grants from BBSRC, Wellcome and the University of Manchester Strategic Fund. The funders had no role in study design, data collection and analysis, decision to publish, or preparation of the manuscript. For the purpose of Open Access, the author has applied a CC BY public copyright licence to any Author Accepted Manuscript version arising from this submission.

## Author contributions

Conceptualisation, M.B and M.A.D-C; methodology, R.K.G, M.B and M.A.D-C; investigation, K.A.S, T.T.M, C.T, S.T, R.F-G, C.B, S.C and M.A.D-C; data curation, K.A.S, T.T.M and M.A.D-C; formal analysis, K.A.S, A.A and M.A.D-C; resources, T.L, R.K.G, P.N, M.B and M.A.D-C; funding acquisition, R.K.G, M.B and M.A.D-C; supervision, C.C, M.B and M.A.D-C; validation, K.A.S and M.A.D-C; visualization, K.A.S, A.A and M.A.D-C; writing of the original draft, K.A.S, M. B and M.A.D-C; writing–review and editing, K.A.S, A.A, R.K.G, P.N, M. B and M.A.D-C.

## Data and code availability

The metagenomic datasets generated during and analysed in the current study are available in the European Nucleotide Archive (ENA) repository (https://www.ebi.ac.uk/ena/browser/home) under the project accession number PRXXXXXXX. Sample metadata is available in *Supplementary Table 17*. The custom pipeline used for downloading and quality-filtering the caecal metagenomic samples can be found as a snakemake pipeline ^73^ in: https://github.com/alexmsalmeida/metagen-fetch.

The R code used to analyse the metagenomics data of this study is available in the Github repository: https://github.com/Duque-Correa-Lab/

---

**Supplementary Figure 1.**
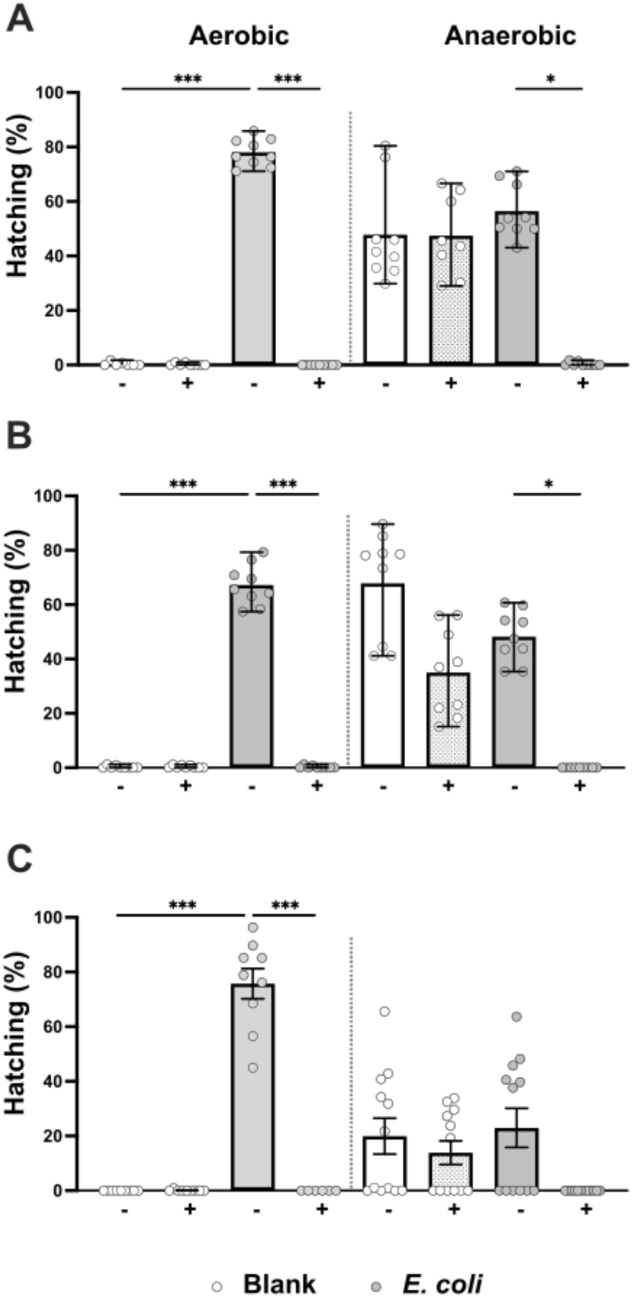
*In vitro* hatching of *Trichuris muris* eggs in response to Escherichia coli. *Trichuris muris* eggs were co-cultured with RPMI-1640 media (blank) and *Escherichia coli* in the absence (-) or presence (+) of a protease inhibitor cocktail (2x) under aerobic and anaerobic conditions at 37°C for 24 h. The number of total embryonated eggs and hatched larvae were counted, from which the hatching percentage was calculated. Experiments were completed in triplicate across 3-4 independent experiments, as negative (blank) and positive (*E. coli*) controls for hatching assays of *T. muris* eggs in co-culture with caecal luminal contents and mucosal scrapings from (A) wild-type (WT), (B) Donor 2 and (C) Donor 7 human-microbiota associated (HMA) mice presented in Figure 4. Each dot represents a well of eggs (n = 7-12). Median and interquartile range are shown, and statistical differences between groups were evaluated using Kruskal Wallis with Dunn’s multiple comparison tests (\**p*<0.05, \*\*\**p*<0.0001).

**Supplementary Figure 2.**
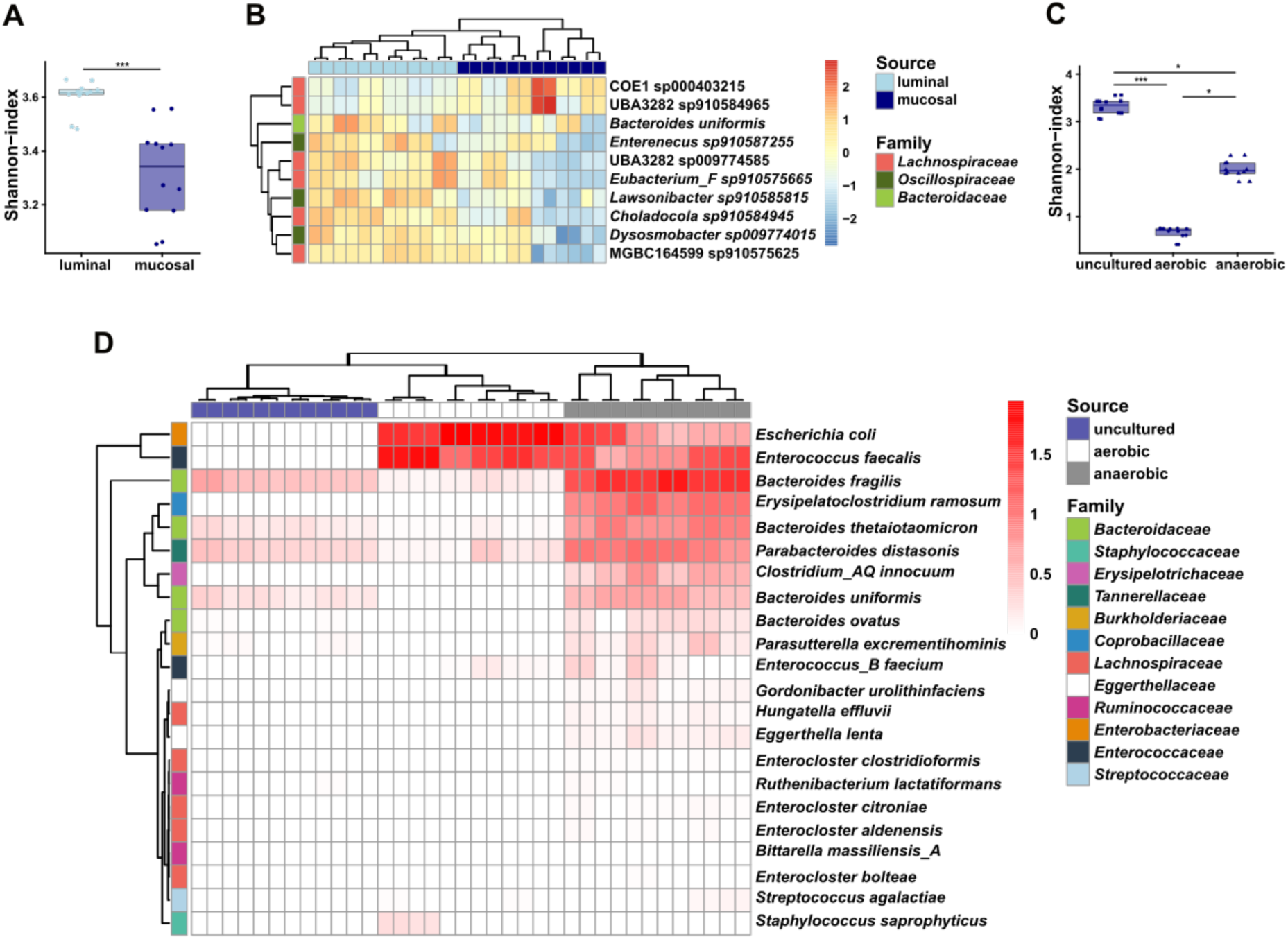
Identification of bacterial species with increased abundance within the mucosal niche of the caecum of Donor 2 human microbiota-associated mice, which are associated with *Trichuris trichiura* hatching. (A and B) Caecal microbial community structure at species level of the luminal content (luminal), and mucosal scraping (mucosal) of Donor 2 (D2) human microbiota-associated (HMA) mice (n = 6 mice, with sequencing done in duplicate). (A) Alpha-diversity (Shannon) index. Median and interquartile range are shown. Statistical differences between groups were evaluated using Kruskal Wallis with Dunn’s multiple comparison tests (\*\*\**p*<0.0001). (B) Heatmap depicting the relative abundance of the ten most abundant microbial species that show statistically significant differences between the lumen (luminal content) and mucosa (mucosal scraping) of the caecum in D2 HMA mice. The heatmap uses a colour scale to represent relative abundances, highlighting variations across the two microbial niches. Clustering was performed using hierarchical clustering with complete linkage on Euclidean dissimilarity matrices. Each column represents a microbial sample. (C and D) Microbial community structure at species level of the caecal mucosa of D2 HMA mice before (uncultured) and after 24 hours of culture under aerobic or anaerobic conditions (n = 6 mice, with sequencing done in duplicate). (C) Alpha-diversity (Shannon) index. Median and interquartile range are shown. Statistical differences between groups were evaluated using Kruskal Wallis test. (\**p*<0.05, \*\*\**p*<0.0001). (D) Heatmap depicting the abundance of the microbial species significantly more abundant in the aerobic and anaerobic metascrapes compared to the uncultured samples of the caecal mucosal scrapings of D2 HMA mice. The heatmap uses a colour scale to represent relative abundances, highlighting variations across the two culture conditions. Clustering was performed using hierarchical clustering with complete linkage on Euclidean dissimilarity matrices. Each column represents a microbial sample.

**Supplementary Figure 3.**
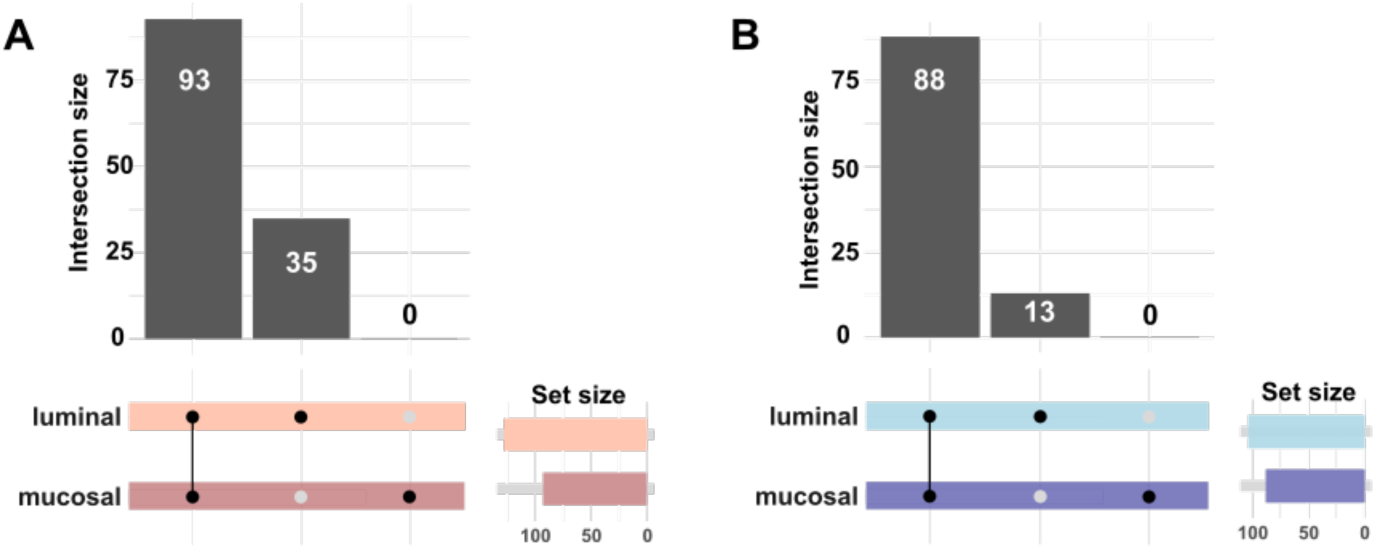
The caecal mucosal microbiome is a subset of the bacteria populations present in the caecal luminal content of human microbiota-associated mice. Upset plot showing shared and unique bacterial species between the luminal content (luminal) and mucosal scraping (mucosal) of (A) Donor 7 and (B) Donor 2 human microbiota-associated mice. Vertical bar plots represent the number of species found in the sample group highlighted in the lower panel. Horizontal bar plots on the right-side panel show the total number of species detected in each mucosal niche.

**Supplementary Figure 4.**
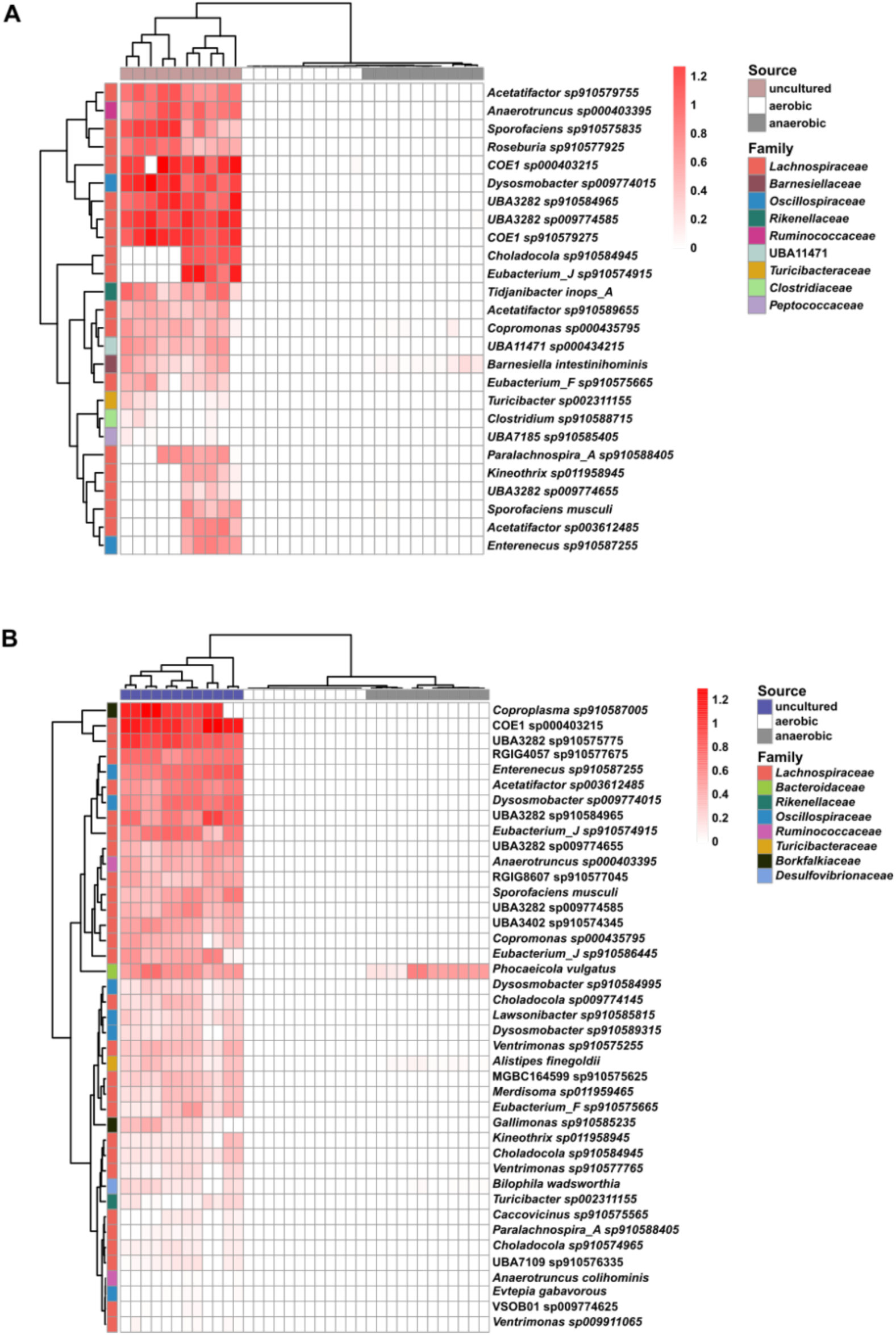
Bacterial species of high abundance in the uncultured caecal mucosal scrapings of Donor 7 and Donor 2 human microbiota-associated mice that are lost upon aerobic and anaerobic culture. Heatmap depicting the relative abundance of all the microbial species statistically significantly more abundant in the uncultured caecal mucosal scrapings compared to the aerobic and anaerobic mucosal metascrapes of (A) Donor 7 and (B) Donor 2 human microbiome-associated mice. The heatmap uses a colour scale to represent relative abundances, highlighting variations across the culture conditions. Clustering was performed using hierarchical clustering with complete linkage on Euclidean dissimilarity matrices. Each column represents a microbial sample.

**Supplementary Figure 5.**
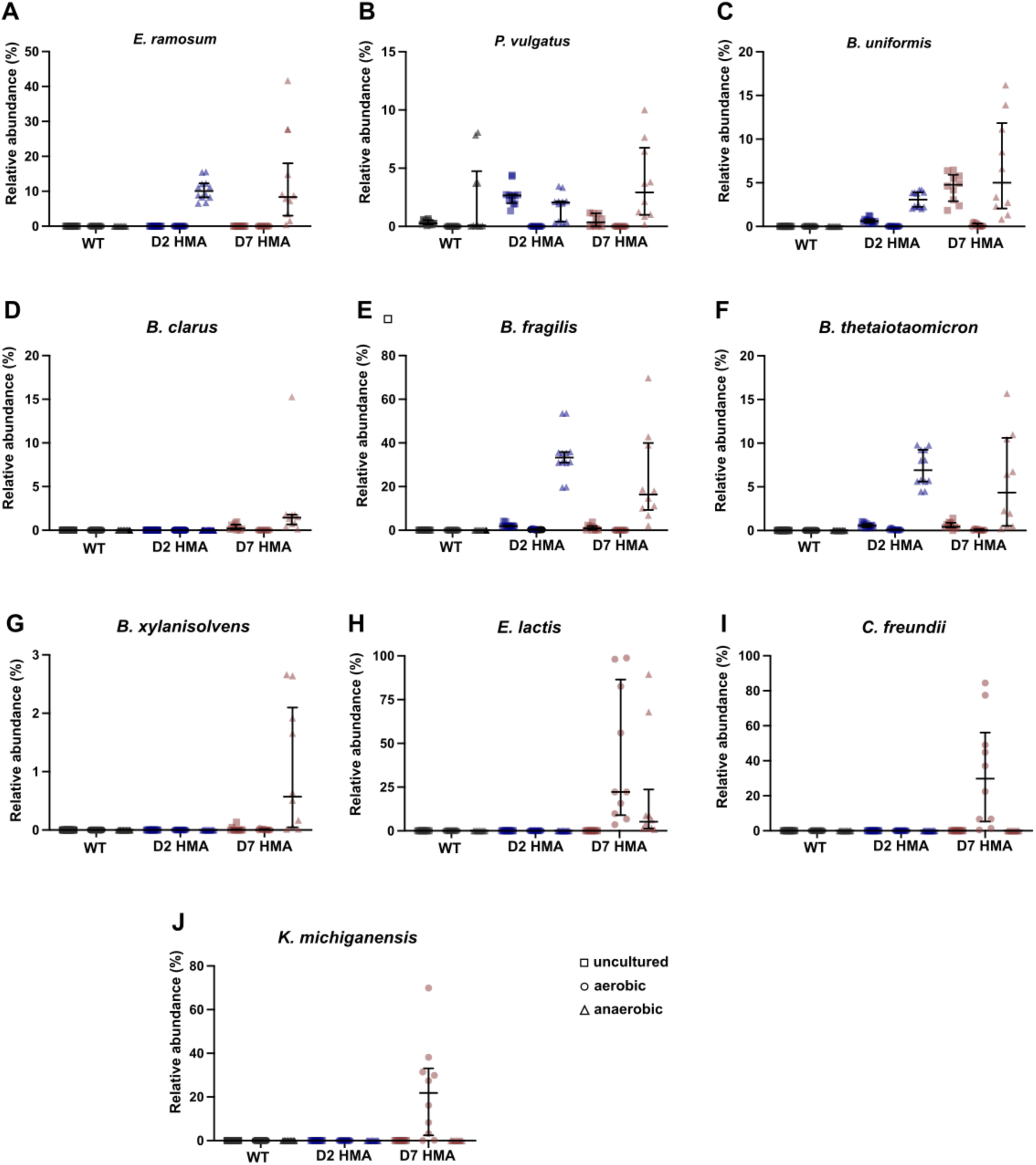
Relative abundances across mouse lines of bacterial species overgrowing in the aerobic and anaerobic metascrapes of the caecal mucosa of Donor 7 human microbiota-associated mice. Relative abundances of *Erysipelatoclostridium ramosum* (A), *Phocaeicola vulgatus* (B), *Bacteroides uniformis* (C), *Bacteroides clarus* (D)*, Bacteroides fragilis* (E)*, Bacteroides thetaiotaomicron* (F), *Bacteroides xylanisolvens* (G), *Enterococcus B lactis* (H)*, Citrobacter freundii* (I), and *Klebsiella michiganensis* (J) in the caecal mucosal scrapings of wild-type (WT), donor 2 (D2) and donor 7 (D7) human microbiota-associated (HMA) mice. Samples were uncultured (uncultured) or cultured for 24 hours under aerobic or anaerobic conditions and DNA from metascrapes was sequenced (WT n = 6 mice with sequencing done in duplicate, D2 HMA n = 6 mice with sequencing done in duplicate, D7 HMA n = 10 mice). Median and interquartile range are shown.

**Supplementary Figure 6.**
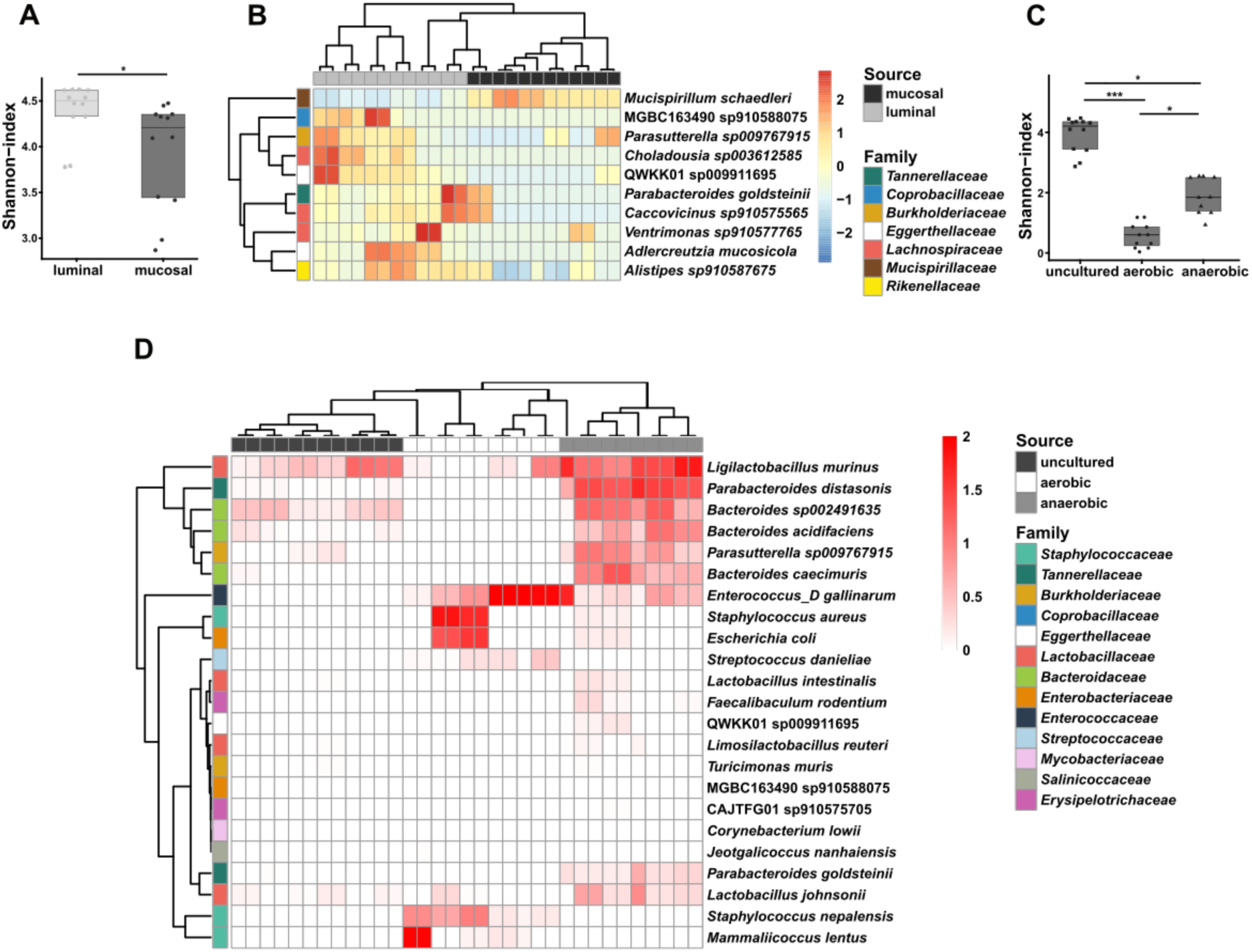
Microbial community structure of the caecal luminal contents and mucosal scrapings of wild-type mice. (A and B) Caecal microbial community structure at species level of the luminal content (luminal), and mucosal scraping (mucosal) of wild-type (WT) mice (n = 6 mice, with sequencing done in duplicate). (A) Alpha-diversity (Shannon) index. Median and interquartile range are shown, and statistical differences between groups were evaluated using Kruskal Wallis with Dunn’s multiple comparison tests (\**p*<0.05). (B) Heatmap depicting the relative abundance of the ten most abundant microbial species that show statistically significant differences between the lumen (luminal content) and mucosa (mucosal scraping) of the caecum in WT mice. The heatmap uses a colour scale to represent relative abundances, highlighting variations across the two microbial niches. Clustering was performed using hierarchical clustering with complete linkage on Euclidean dissimilarity matrices. Each column represents a microbial sample. (C and D) Microbial community structure at species level of the caecal mucosa of WT mice before (uncultured) and after 24 hours of culture under aerobic or anaerobic conditions (n = 6 mice, with sequencing done in duplicate). (C) Alpha-diversity (Shannon) index. Median and interquartile range are shown, and statistical differences between groups were evaluated using Kruskal Wallis test (\**p*<0.05, \*\*\**p*<0.001). (D) Heatmap depicting the abundance of the microbial species significantly more abundant in the aerobic and anaerobic metascrapes compared to the uncultured samples of the caecal mucosal scrapings of WT mice. The heatmap uses a colour scale to represent relative abundances, highlighting variations across the two culture conditions. Clustering was performed using hierarchical clustering with complete linkage on Euclidean dissimilarity matrices. Each column represents a microbial sample.

## Supplementary Table Legends

**Supplementary Table 1. Abundances of microbial taxa at species level in the caecal microbiome of wild-type and human microbiota-associated mice.**

Abundances of microbial taxa at species level found in the whole caecum of wild-type (WT, n = 3), Donor 2 (D2, n = 3), and Donor 7 (D7, n = 4) human microbiota-associated (HMA) mice. Bacterial abundance data were generated using MetaPhlAn 4 (v4.0.240) applying a minimal prevalence threshold of 0.01.

**Supplementary Table 2. Unique and shared bacterial species in the caecal microbiome of wild-type and human microbiota-associated mice.**

Absence (0) or presence (1) of bacterial species in the caecal microbiome of wild-type (WT, n = 3), Donor 2 (D2, n = 3) and Donor 7 (D7, n = 4) human microbiota-associated (HMA) mice. A bacterial species was considered present if its abundance was greater than zero in at least one sample within the respective group.

**Supplementary Table 3. Bacterial species with statistically significant abundance differences between wild-type and human microbiota-associated mice.**

Caecal microbial samples from wild-type (WT), Donor 2 (D2) and Donor 7 (D7) human microbiota-associated (HMA) mice were collected and differentially abundant bacterial species identified. Data was obtained using Maaslin2 function from the MaAsLin2 package in R for performing multivariate association analysis. Log transformation was performed on the data, and no normalisation applied. The effect size indicates the direction and magnitude of association between the metadata variable (experimental group) and the microbial feature (species abundance). Negative and positive effect size values indicated lower and higher species abundance in the caecum of the particular experimental group (WT or D2 HMA) in comparison to D7 HMA, respectively. The standard error of the effect size (stderr) quantifies the uncertainty of the estimate. A species needed to be present in at least 1% of samples to be included in the analysis, and only associations with an adjusted *p*-value (q) ≤ 0.05 were considered significant. The analysis corrects for multiple testing using the Benjamini-Hochberg method. Species unique to the D7 HMA mouse microbiome are highlighted.

**Supplementary Table 4. Abundances of microbial taxa at species level in the caecal microbiome of Donor 7 human microbiota-associated mice.**

Abundances of microbial taxa at species level found in the caecal luminal contents and mucosal scrapings of Donor 7 (D7) human microbiota-associated (HMA) mice (n = 10). Bacterial abundances were profiled using MetaPhlAn 4 (v4.0.240) applying a minimum prevalence threshold of 0.01. Samples are grouped by individual mouse, with each entry annotated by a unique ‘Sample title’ corresponding to those shown in Supplementary Table 17.

**Supplementary Table 5. Abundances of microbial taxa at species level in the caecal microbiome of Donor 2 human microbiota-associated mice.**

Abundances of microbial taxa at species level found in the caecal luminal contents and mucosal scrapings of Donor 2 (D2) human microbiota-associated (HMA) mice (n = 6 mice, with sequencing done in duplicate). Bacterial abundances were profiled using MetaPhlAn 4 (v4.0.240) applying a minimum prevalence threshold of 0.01. Samples are grouped by individual mouse, with each entry annotated by a unique ‘Sample title’ corresponding to those shown in Supplementary Table 17.

**Supplementary Table 6. Bacterial species with significant abundance differences between the caecal mucosal and luminal microbiome of the Donor 7 human microbiota-associated mice.**

Caecal microbial samples from the luminal contents or mucosal scrapings of the Donor 7 (D7) human microbiota-associated (HMA) mice were collected and differentially abundant bacterial species identified. Data was obtained using Maaslin2 function from the MaAsLin2 package in R for performing multivariate association analysis. Log transformation was performed on the data, and no normalisation applied. The effect size indicates the direction and magnitude of association between the metadata variable (experimental group) and the microbial feature (species abundance). Negative and positive effect size values indicated lower and higher species abundance of bacterial species in the lumen in comparison to the mucosa. The standard error of the effect size (stderr) quantifies the uncertainty of the estimate. A species needed to be present in at least 1% of samples to be included in the analysis, and only associations with an adjusted *p*-value (q) ≤ 0.05 were considered significant. The analysis corrects for multiple testing using the Benjamini-Hochberg method.

**Supplementary Table 7. Bacterial species with significant abundance differences between the caecal mucosal and luminal microbiome of the Donor 2 human microbiota-associated mice.**

Caecal microbial samples from the luminal contents or mucosal scrapings of the Donor 2 (D2) human microbiota-associated (HMA) mice were collected and differentially abundant bacterial species identified. Data was obtained using Maaslin2 function from the MaAsLin2 package in R for performing multivariate association analysis. Log transformation was performed on the data, and no normalisation applied. The effect size indicates the direction and magnitude of association between the metadata variable (experimental group) and the microbial feature (species abundance). Negative and positive effect size values indicated lower and higher species abundance of bacterial species in the lumen in comparison to the mucosa. The standard error of the effect size (stderr) quantifies the uncertainty of the estimate. A species needed to be present in at least 1% of samples to be included in the analysis, and only associations with an adjusted *p*-value (q) ≤ 0.05 were considered significant. The analysis corrects for multiple testing using the Benjamini-Hochberg method.

**Supplementary Table 8. Abundances of microbial taxa at species level in the caecal microbiome and its metascrapes of Donor 7 human microbiota-associated mice.**

Abundances of microbial taxa at species level found in the mucosal scraping of Donor 7 (D7) human microbiota-associated (HMA) mice (n = 10), and its metascrapes achieved by culturing mucosal scraping samples for 24 hours under either aerobic or anaerobic conditions. Bacterial abundances were profiled using MetaPhlAn 4 (v4.0.240) applying a minimum prevalence threshold of 0.01. Samples are grouped by individual mouse, with each entry annotated by a unique ‘Sample title’ corresponding to those shown in Supplementary Table 17.

**Supplementary Table 9. Abundances of microbial taxa at species level in the caecal microbiome and its metascrapes of Donor 2 human microbiota-associated mice.**

Abundances of microbial taxa at species level found in the mucosal scraping of Donor 2 (D2) human microbiota-associated (HMA) mice (n = 6 with sequencing done in duplicate), and its metascrapes achieved by culturing mucosal scraping samples for 24 hours under either aerobic or anaerobic conditions. Bacterial abundances were profiled using MetaPhlAn 4 (v4.0.240) applying a minimum prevalence threshold of 0.01. Samples are grouped by individual mouse, with each entry annotated by a unique ‘Sample title’ corresponding to those shown in Supplementary Table 17.

**Supplementary Table 10. Bacterial species with significant abundance differences between the metascrapes and uncultured microbiome of the caecal mucosal scraping of Donor 7 human microbiota-associated mice.**

Microbial samples from the caecal mucosa of Donor 7 (D7) human microbiota-associated (HMA) mice and its aerobic and anaerobic metascrapes were collected, and differentially abundant bacterial species identified. Data was obtained using Maaslin2 function from the MaAsLin2 package in R for performing multivariate association analysis. Log transformation was performed on the data, and no normalisation applied. The effect size indicates the direction and magnitude of association between the metadata variable (experimental group) and the microbial feature (species abundance). Negative and positive effect size values indicated lower and higher species abundance in the metascrapes cultured under the particular condition (aerobic or anaerobic) in comparison to uncultured mucosal scrapings, respectively. The standard error of the effect size (stderr) quantifies the uncertainty of the estimate. A species needed to be present in at least 1% of samples to be included in the analysis, and only associations with an adjusted *p*-value (q) ≤ 0.05 were considered significant. The analysis corrects for multiple testing using the Benjamini-Hochberg method.

**Supplementary Table 11. Bacterial species with significant abundance differences between the metascrapes and uncultured microbiome of the caecal mucosal scraping of Donor 2 human microbiota-associated mice.**

Microbial samples from the caecal mucosa of Donor 2 (D2) human microbiota-associated (HMA) mice and its aerobic and anaerobic metascrapes were collected, and differentially abundant bacterial species identified. Data was obtained using Maaslin2 function from the MaAsLin2 package in R for performing multivariate association analysis. Log transformation was performed on the data, and no normalisation applied. The effect size indicates the direction and magnitude of association between the metadata variable (experimental group) and the microbial feature (species abundance). Negative and positive effect size values indicated lower and higher species abundance in the metascrapes cultured under the particular condition (aerobic or anaerobic) in comparison to uncultured mucosal scrapings, respectively. The standard error of the effect size (stderr) quantifies the uncertainty of the estimate. A species needed to be present in at least 1% of samples to be included in the analysis, and only associations with an adjusted *p*-value (q) ≤ 0.05 were considered significant. The analysis corrects for multiple testing using the Benjamini-Hochberg method.

**Supplementary Table 12. Abundances of microbial taxa at species level in the caecal microbiome and its metascrapes of wild-type mice.**

Abundances of microbial taxa at species level found in the mucosal scraping of wild-type (WT) mice (n = 6 with sequencing done in duplicate), and its metascrapes achieved by culturing mucosal scraping samples for 24 hours under either aerobic or anaerobic conditions. Bacterial abundances were profiled using MetaPhlAn 4 (v4.0.240) applying a minimum prevalence threshold of 0.01. Samples are grouped by individual mouse, with each entry annotated by a unique ‘Sample title’ corresponding to those shown in Supplementary Table 17.

**Supplementary Table 13. *Trichuris trichiura* egg hatching is not triggered *in vitro* by aerobic bacteria present in the caecal mucosa of humanised-microbiota mice.**

*Trichuris trichiura* eggs were co-cultured with Luria Bertani media (blank), or *Klebsiella michiganensis* and *Citrobacter freundii* at 37°C for one, three and six days under aerobic conditions. The number of total embryonated eggs and hatched larvae were counted, from which the percentage of hatching was calculated. Hatching was completed in triplicate across 3 independent experiments and each dot presents a well of eggs (n = 9).

**Supplementary Table 14. Abundances of microbial taxa at species level in the caecal microbiome of wild-type mice.**

Abundances of microbial taxa at species level found in the caecal luminal content and mucosal scraping of wild-type (WT) mice (n = 6 mice, with sequencing done in duplicate). Bacterial abundances were profiled using MetaPhlAn 4 (v4.0.240) applying a minimum prevalence threshold of 0.01. Samples are grouped by individual mouse, with each entry annotated by a unique ‘Sample title’ corresponding to those shown in Supplementary Table 17.

**Supplementary Table 15. Bacterial species with significant abundance differences between the caecal mucosal and luminal microbiome of wild-type mice.**

Caecal microbial samples from the luminal contents or mucosal scrapings of wild-type (WT) mice were collected and differentially abundant bacterial species identified. Data was obtained using Maaslin2 function from the MaAsLin2 package in R for performing multivariate association analysis. Log transformation was performed on the data, and no normalisation applied. The effect size indicates the direction and magnitude of association between the metadata variable (experimental group) and the microbial feature (species abundance). Negative and positive effect size values indicated lower and higher species abundance of bacterial species in the lumen in comparison to the mucosa. The standard error of the effect size (stderr) quantifies the uncertainty of the estimate. A species needed to be present in at least 1% of samples to be included in the analysis, and only associations with an adjusted *p*-value (q) ≤ 0.05 were considered significant. The analysis corrects for multiple testing using the Benjamini-Hochberg method.

**Supplementary Table 16. Bacterial species with significant abundance differences between the metascrapes and uncultured microbiome of the caecal mucosal scraping of wild-type mice.**

Microbial samples from the caecal mucosa of wild-type (WT) mice and its aerobic and anaerobic metascrapes were collected, and differentially abundant bacterial species identified. Data was obtained using Maaslin2 function from the MaAsLin2 package in R for performing multivariate association analysis. Log transformation was performed on the data, and no normalisation applied. The effect size indicates the direction and magnitude of association between the metadata variable (experimental group) and the microbial feature (species abundance). Negative and positive effect size values indicated lower and higher species abundance in the metascrapes cultured under the particular condition (aerobic or anaerobic) in comparison to uncultured mucosal scrapings, respectively. The standard error of the effect size (stderr) quantifies the uncertainty of the estimate. A species needed to be present in at least 1% of samples to be included in the analysis, and only associations with an adjusted p-value (q) ≤ 0.05 were considered significant. The analysis corrects for multiple testing using the Benjamini-Hochberg method.

**Supplementary Table 17. Sample metadata for metagenomic libraries.** Metagenomic libraries metadata including sample and run accession number, and sample title as found on European Nucleotide Archive (ENA).

